# Using a Deep Generation Network Reveals Neuroanatomical Specificity in Hemispheres

**DOI:** 10.1101/2023.08.18.553830

**Authors:** Gongshu Wang, Ning Jiang, Yunxiao Ma, Tianyi Yan

**Affiliations:** School of Medical Technology, Beijing Institute of Technology, 100081 Beijing, China

## Abstract

Asymmetry is an important property of brain organization, but its nature is still poorly understood. Capturing the neuroanatomical components specific to each hemisphere facilitates the understanding of the establishment of brain asymmetry. Since deep generative networks (DGNs) have powerful inference and recovery capabilities, we use one hemisphere to predict the opposite hemisphere by training the DGNs, which automatically fit the built-in dependencies between the left and right hemispheres. After training, the reconstructed images approximate the homologous components in the hemisphere. We use the difference between the actual and reconstructed hemispheres to measure hemisphere-specific components due to asymmetric expression of environmental and genetic factors. The results show that our model is biologically plausible and that our proposed metric of hemispheric specialization is reliable, representing a wide range of individual variation. Together, this work provides promising tools for exploring brain asymmetry and new insights into self-supervised DGNs for representing the brain.

## Introduction

Although the left (LH) and right (RH) hemispheres of our brains develop with a high degree of symmetry at both the anatomical and functional levels, it has become clear that the two sides develop and age at different rates, have subtle structural and functional differences and are each dominant in processing specific cognitive and perceptual tasks [1–4]. A complex and ongoing interplay of genetic and environmental factors most likely underlies and guides this lateralized specialization. Genes provide templates for creating brain patterns that probably trigger only modest hemispheric dominance, while prenatal and postnatal environments shape and influence the orientation of emerging neural networks, reinforcing hemispheric dominance [5–7].

Understanding the nature of brain asymmetry and how it contributes to individual variation is an enduring crucial topic in neuroscience. However, the methods currently in use for addressing these problems remain unsatisfactory. For example, calculating the asymmetric index [8–10] by comparing the LH and RH, but it only indicates the degree of lateralization without revealing the neuroanatomical details responsible for the lateralization.

According to brain development and lateralization [1, 7], the process of lateralized specialization can be considered the coupling of shared factors and unique factors. Shared factors (such as most genes) underlie the formation of both the LH and RH, while unique factors (such as asymmetric light, posture and injury) dominate the direction of development of only one hemisphere. As a result, in real-world scenarios, the LH and RH are structured by homologous components and hemisphere-specific components (Fig. 1a). We believe that uncovering hemisphere-specific components will provide deeper insights into brain asymmetries. However, with limited a priori knowledge, it is difficult to disentangle these two components directly.

**Fig. 1.**
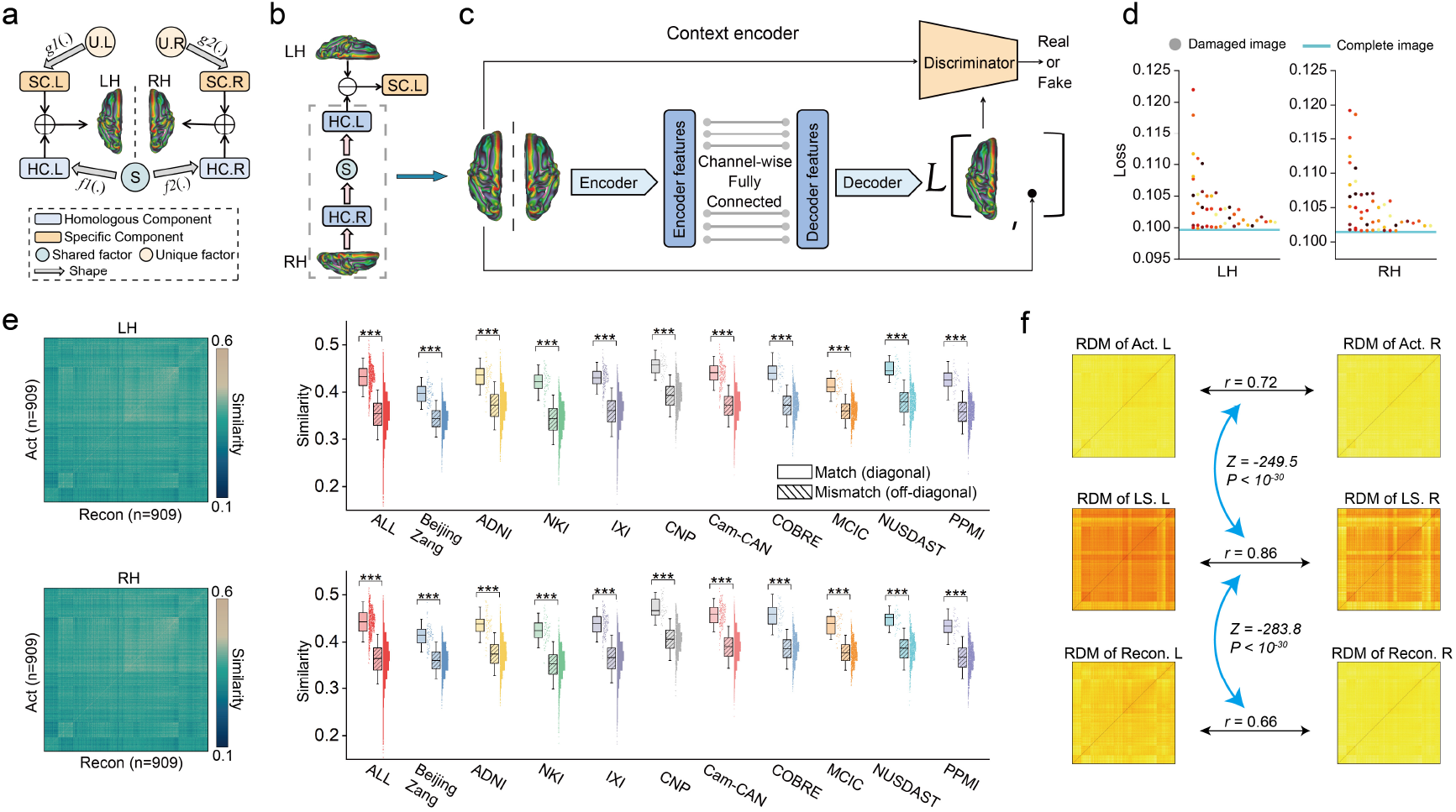
Train the DGNs to use one hemisphere to predict the opposite hemisphere. **(a)** Brain hemispheres are composed of homologous and specific components. **(b)** Using CE to establish a mapping between homologous components of the LH and RH, which follows the principle of hemisphere formation. **(c)** The CE Architecture. The generator consists of a CNN encoder, a channelwise fully connected layer and a CNN decoder. The discriminator also consists of a stack of CNN layers. We trained two CE models to implement the prediction of LH to RH and RH to LH. **(d)** Averaged reconstruction loss across subjects when the model is fed with ‘virtual lesion’ images. Each point represents a class of damaged images (n = 58). The blue line indicates reconstruction loss when the model is fed with complete images. **(e)** The similarity between all pairs of reconstructed and actual hemisphere images. In the heatmaps, diagonal values (n = 909) indicate similarity between matched images, and off-diagonal values (n = 825372) indicate similarity between unmatched images. The data in the box range from 25% to 75%. Error bars indicate standard deviation (SD) ×1.5. *** indicates p < 10^-30^, using a two-sample t test. **(f)** RSA between LH and RH. Black arrows indicate the Spearman correlation analysis between RDMs. Blue arrows indicate comparisons between correlation coefficients using the Steiger’s z test. HC: homologous component, SC: specific component, S: shared factor, U: unique factor. *f1*(.) and *f2*(.) denote the mapping from S to HC. *g1*(.) and *g2*(.) denote the mapping from U to SC. ‘Act’ indicates the actual hemisphere, ‘Recon’ indicates the reconstructed hemisphere, and ‘LS’ indicates the latent space (feature map in the bottleneck of the model).

Assuming an ideal situation in which the LH and RH are shaped only by shared factors, it is possible to establish an unambiguous mapping between them because they consist only of strongly dependent homologous components (similar to homotopy invariants in topology) [11, 12]. When this ideal mapping function is applied to the real world, we can leverage the actual LH (resp. RH) to reconstruct a “fake” RH (resp. LH) consisting of homologous components. Afterward, by comparing the actual hemispheres with the ‘fake’ hemispheres, we can infer hemisphere-specific components derived from the asymmetrical influence of unique factors. Here, we aimed to uncover the connections between homologous components and later reveal hemispheric specializations.

Deep generative networks (DGNs) provide reliable tools for image composition, which can accurately perform style transformations [13], such as transforming facial photos into cartoon styles. Currently, they have been widely applied in the synthesis of high-quality brain images [14, 15]. In addition, DGNs have the powerful ability to learn to recover images by viewing corrupted examples without ground truth data training. For instance, in the absence of clean samples, they can achieve effective denoising by building a mapping between pairs of noisy images [16] or by self-supervised learning on a single noisy image[17]. In the LH and RH, their homologous components are strongly dependent, while their specific components are relatively independent. Thus, even without ground truth data (e.g., when the brain is shaped only by shared factors), it is possible to capture homologous components from empirical neuroimages by training DGNs.

In pursuit of both high-quality generated images and biologically reliable mapping, we modified a state-of-the-art DGN, the context encoder (CE), to implement complex nonlinear transformations. CEs are unsupervised learning algorithms that can generate the content of an arbitrary image region conditioned on its surrounding environment [18]. They are jointly trained by minimizing both a reconstruction loss and an adversarial loss [19], which can produce much sharper results. CE-inspired methods are conducive to capturing biological representations from medical imaging [20, 21]. When CE is applied to brain magnetic resonance imaging (MRI), the generated images are more likely to have a biologically semantic structure rather than just a visual appearance.

In this study, we used a large sample of gray matter (GM) maps (n = 2852), an indirect measure of the complex structure of neurons, to train and validate the models. The goal of the models was to convert one side of the hemisphere to fit the opposite hemisphere as well as possible. During training, to find the optimal solution, the models continuously approximated inherent dependent patterns between homologous components. The computational process essentially follows the principle of brain lateralization, which focuses on meaningful features, encodes them into a common latent space (similar to shared factors), and later decodes them in another space (Fig.1b).

We trained one CE to implement the conversion from LH to RH and another CE to implement conversion from RH to LH. We used the difference between the reconstructed and actual hemispheres as a biological indicator to estimate the neuroanatomical specialization in the hemisphere. Evidence has suggested that model failures can be informative[22]. Age-prediction error[23], for example, is of value for assessing normal aging and disease[24, 25]. This provides strong support for the reliability of our proposed biological indicators.

First, we verified that the trained models are biologically plausible and that our proposed hemispheric-specialization features are stable and reliable. Second, we demonstrated that hemispheric specialization features are important biological indicators that are differentially related to age, sex, disease, environment, and cognition. Then, we demonstrated that the spatial distribution of hemispheric specialization features is plausible and found that LH and RH have unequal contributions to hemispheric specialization. In addition, through comprehensive analysis, we identified some crucial regions that account for most of the variation in hemispheric specialization and that control and drive the hemispheric specialization process.

## Results

### The reconstructed hemispheres are biologically plausible

We used 1372 healthy subjects from ten datasets (Table 1) to train two CE models (Fig. 1c) and verified the biological plausibility of the model from several aspects.

**Table 1.**
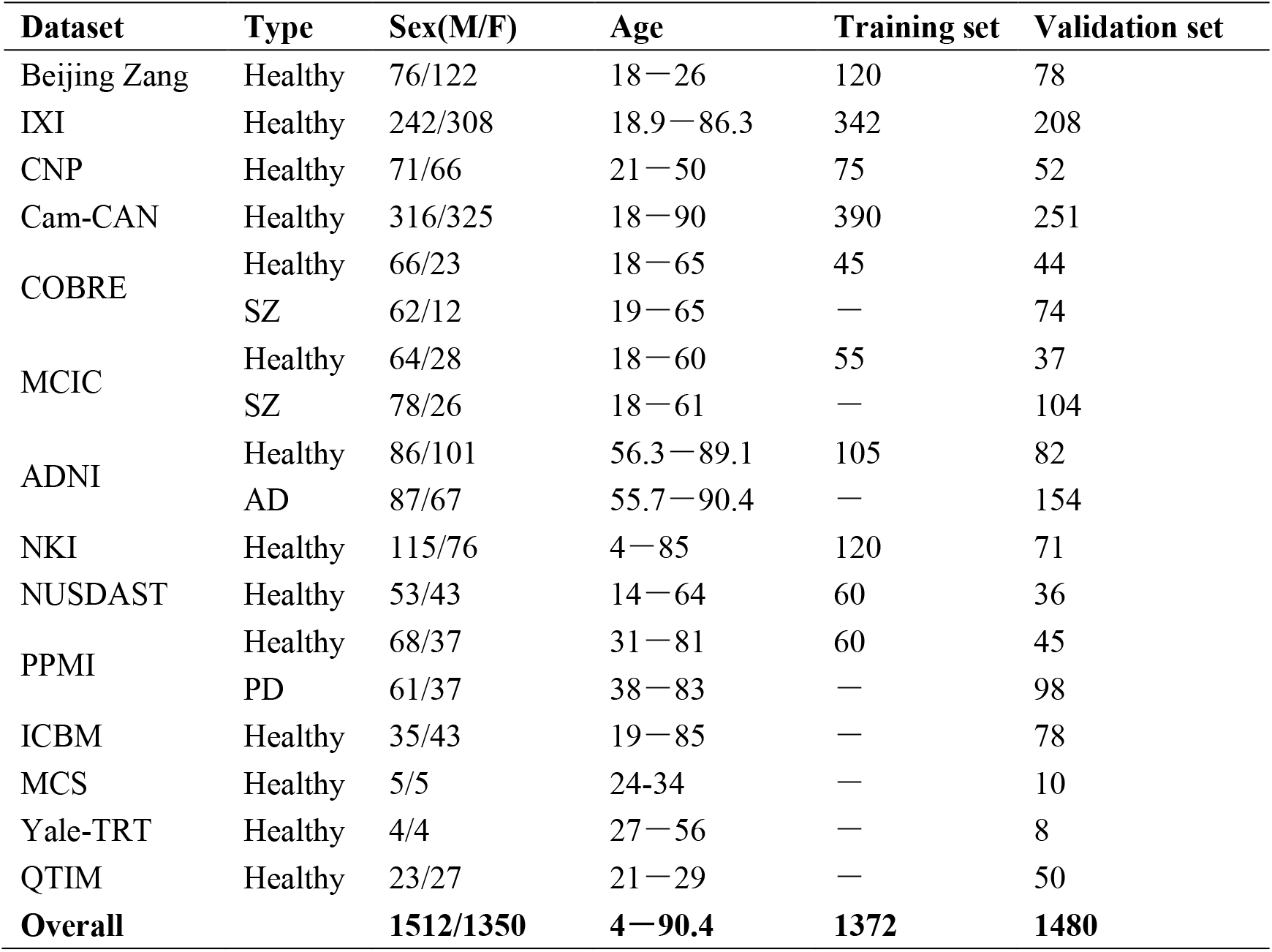
Information about the datasets.

| Dataset | Type | Sex(M/F) | Age | Training set | Validation set |
| --- | --- | --- | --- | --- | --- |
| Beijing Zang | Healthy | 76/122 | 18—26 | 120 | 78 |
| IXI | Healthy | 242/308 | 18.9—86.3 | 342 | 208 |
| CNP | Healthy | 71/66 | 21—50 | 75 | 52 |
| Cam-CAN | Healthy | 316/325 | 18—90 | 390 | 251 |
| COBRE | Healthy | 66/23 | 18—65 | 45 | 44 |
|  | SZ | 62/12 | 19—65 | — | 74 |
| MCIC | Healthy | 64/28 | 18—60 | 55 | 37 |
|  | SZ | 78/26 | 18—61 | — | 104 |
| ADNI | Healthy | 86/101 | 56.3—89.1 | 105 | 82 |
|  | AD | 87/67 | 55.7—90.4 | — | 154 |
| NKI | Healthy | 115/76 | 4—85 | 120 | 71 |
| NUSDAST | Healthy | 53/43 | 14—64 | 60 | 36 |
| PPMI | Healthy | 68/37 | 31—81 | 60 | 45 |
|  | PD | 61/37 | 38—83 | — | 98 |
| ICBM | Healthy | 35/43 | 19—85 | — | 78 |
| MCS | Healthy | 5/5 | 24-34 | — | 10 |
| Yale-TRT | Healthy | 4/4 | 27—56 | — | 8 |
| QTIM | Healthy | 23/27 | 21—29 | — | 50 |
| **Overall** |  | **1512/1350** | **4—90.4** | **1372** | **1480** |

First, we estimated whether the CE-based models were applicable to the brain MRI data. We iteratively fed the ‘virtual lesion’ hemispheric images, in which a region is masked, into the trained model and then calculated the reconstruction loss. We found that the loss values of the virtual lesion images were all higher than the loss values of the complete images (Fig. 1d), indicating that each brain region contributes positively to the generation of images. This result suggests that the model focused on the whole brain and did not neglect information in any region. In addition, we used an independent dataset ICBM to evaluate the optimization quality during training. As the number of training epochs increases, the quality of the reconstructed images improves in both the training set and the independent validation set. Thus, the models are not overfitted and are robust across data domains.

Then, we examined whether the CE models captured the semantic information in the brain. We calculated the pairwise similarity between all pairs of actual and reconstructed images (heatmaps in Fig. 1e). The results showed that within-individual similarity was significantly higher than between-individual similarity both in the combination of all datasets and within each dataset (p values < 10^-30^, two-sample t test, box plots in Fig. 1e). Thus, our models can capture the semantic information related to individual identity and retain it in the reconstructed brain.

Furthermore, we tested whether our model tended to capture the shared factors underlying LH and RH. We constructed the representational dissimilarity matrices (RDMs) using the actual and reconstructed hemispheres as well as the latent space of the models (extracted from the bottleneck of the models). The representational similarity analysis (RSA) showed that the similarity between the actual and reconstructed LH and RH was significantly lower than the similarity between the latent spaces of LH and RH (z = −249.5 and −283.8, p values < 10^-30^, Steiger’s z test, Fig. 1f). In addition, the interindividual similarity increased as the layers became deeper (p = 0, one-way ANOVA, Supplementary Fig. 1). Thus, our models simulated the inverse process of brain evolution, which automatically encoded LH and RH into a relatively shared space.

Collectively, these findings demonstrate the plausibility of our trained CE models, which characterize all brain voxels, capture the idiosyncratic pattern of an individual and follow the principles of brain development and lateralization.

### Calculation of hemisphere-specific neuroanatomical components

Afterward, we used the trained models to calculate the hemispheric specialization of the remaining subjects. To better characterize hemispheric specialization, we applied two metrics: absolute neuroanatomical specificity (ANS) and relative neuroanatomical specialization (RNS), which indicate the amount and proportion, respectively, of specific components in a voxel of LH or RH (see Methods). We visualized the averaged ANS and RNS maps of the LH and RH across 909 healthy subjects from the validation set (Fig. 2a and b, see Supplementary Fig. 2 for a flat surface). There is no difference between the overall ANS and RNS of the LH and RH (Supplementary Fig. 3). To facilitate the analysis of the biological properties of ANS or RNS, we sorted the voxels in LH or RH into 10 equal parts (regions) according to their ANS or RNS values. The mask for each part is shown in Supplementary Fig. 4. The RSA between parts is shown in Supplementary Fig. 5.

**Fig. 2.**
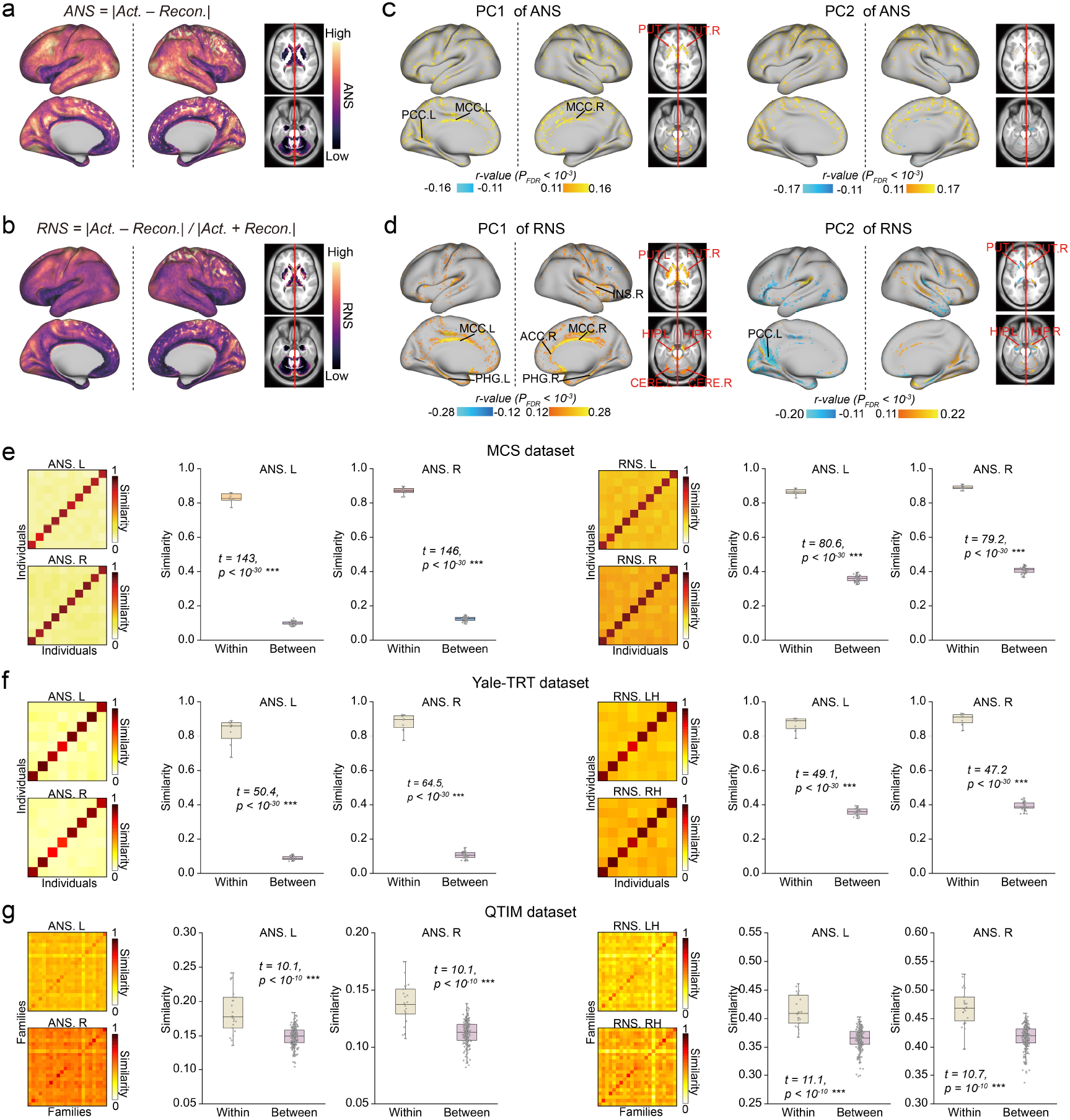
Calculation and validation of hemispheric specialization features. **(a)** and **(b)** Definition of ANS and RNS and visualization of ANS and RNS maps for LH and RH (averaged across healthy people in the validation set, n = 909). **(c)** and **(d)** The main variant loci of ANS and RNS across individuals. Areas most relevant to the first two PCs of ANS and RNS (Pearson partial correlation analysis, thresholds are illustrated in the figure, p values were corrected by FDR). Notably, in (a) – (d), the analyses were performed separately for LH and RH, and we visualized them together. **(e)** and **(f)** Similarity between all paired ANSs or RNSs of the two scans in the MCS dataset and Yale-TRT dataset. Diagonal values indicate similarity within individuals (n = 10 and 8), and off-diagonal values (n = 45 and 28) indicate similarity between individuals. **(g)** The similarity between all pairs of ANSs or RNSs of the two families in the QTIM dataset. Diagonal values indicate similarity within families (n = 22), and off-diagonal values (n = 231) indicate similarity between families. The two-sided two-sample t test was used for between-group comparisons. The data in the box range from 25% to 75%. Error bars indicate SD×1.5. INS: Insula, ACC: Anterior Cingulate Cortex, MCC: Median Cingulate Cortex, PCC: Posterior Cingulate Cortex, PHA: Parahippocampal Area, PUT: Putamen, HIP: Hippocampus, CERE: Cerebellum.

Furthermore, we aimed to identify loci of variation in hemispheric ANS/RNS among individuals. To represent the major variation, we computed the first two principal components (PCs) of the hemispheric ANS/RNS maps across 909 healthy subjects from the validation set. We later calculated the correlation between PCs and ANS/RNS for each voxel (Pearson partial correlation using age, sex, and scanner as covariates, Fig. 2c and d). The results suggest that the important loci of ANS are relatively scattered. In contrast, the important loci of the RNS were relatively concentrated and mainly in the cingulate cortex, subcortex, and cerebellum. Of note, these regions had lower ANS and RNS (Fig. 2a and b). Labeling of regions was performed according to the AAL atlas [26] and Glasser et al. [27]

Next, we verified the reliability of the ANS and RNS values. First, we tested whether ANS and RNS were robust over two scan sessions. We estimated the ANS and RNS of MCS and Yale-TRT individuals and calculated pairwise similarity between all pairs of scan sessions (heatmaps in Fig. 2e and f). We found that within-individual similarity was significantly higher than between-individual similarity for both ANS and RNS of the LH and RH (p values<10^-30^, two-sample t test; box plots in Fig. 2e and f). This result suggests that the ANS and RNS are stable over multiple scans using the same scanner or different scanners.

Second, we tested whether ANS and RNS are heritable in relatives. We estimated the ANS and RNS of QTIM individuals and calculated pairwise similarity between all pairs of families (heatmaps in Fig. 2g). Within-family similarity was significantly higher than between-family similarity (p values<10^-30^, two-sample t test; box plots in Fig. 2g). This result suggests that similar living environments lead to similar ANS and RNS, which is consistent with the conclusion that the degree of brain lateralization is heritable [28].

In addition, we compared our proposed ANS and RNS with the traditional asymmetric index [8]. The results suggest that ANS and RNS are more stable and reliable indicators in identifying interindividual variation, especially in identifying families (Supplementary Fig. 6 and Supplementary Table. 1).

Together, these results suggest that ANS and RNS are biologically meaningful and may be used to represent brain “fingerprinting” and explain variations between individuals.

### ANS and RNS are associated with age and sex

Aging and gender all have a significant impact on brain neuroanatomy and asymmetry [29–31]. In this section, to identify the biological characteristics of the hemispheric specialization features, we analyzed the relationship between the ANS and RNS and these phenotypes.

Brain aging is generally associated with a loss of GM tissue so GM can be used to accurately predict chronological age [32]. We first examined whether the reconstructed brain preserved age-related semantic information. Participants in the training set were used to build dimensionality reduction and regression models, which were later applied to the actual and reconstructed images in the validation set (Fig. 3a). We found no significant difference (p > 0.05, paired samples t test) between the ages predicted from the actual and reconstructed images. Likewise, to demonstrate that the reconstructed images also preserve gender-related information, we used the training set to build a gender classification model, which was then applied to the actual and reconstructed images in the validation set. The classification performance of actual and reconstructed images is highly similar (Supplementary Table. 2). Thus, models have similar perceptions [33] of actual and reconstructed images, indicating that the reconstructed images preserve the inherent patterns of the brain and have the potential to provide meaningful insights into the brain.

**Fig. 3.**
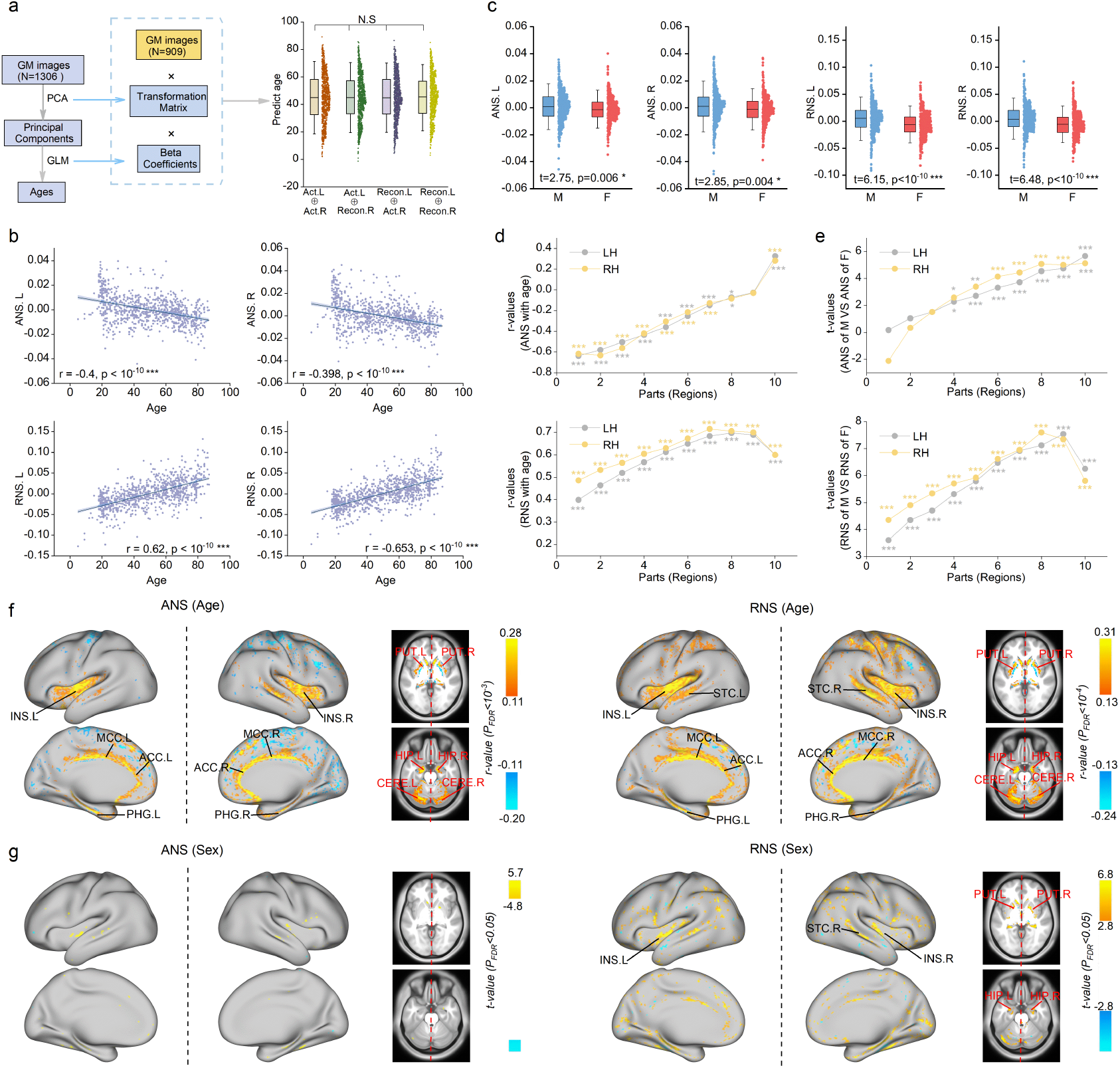
Association between ANS/RNS with age and sex. **(a)** Predicted age (n = 903) based on actual or generated hemisphere images. The transformation matrix of PCA and the beta coefficients of the regression model were estimated from the training set. **(b)** Correlation between mean hemispheric ANS/RNS with age (n = 903, Spearman correlation analysis, since the subjects’ ages did not obey a normal distribution). Gender and scanner information was removed from the ANS/RNS before calculation. The shaded area represents the 95% confidence interval. **(c)** Differences between the mean hemispheric ANS/RNS of men (n=486) and women (n=417). Two-sided two-sample t test. Age and scanner information was removed from the ANS/RNS before calculation. **(d)** and **(e)** Correlation analysis with age and comparison with sex were performed separately in each of the 10 parts (Supplementary Fig. 4). **(f)** The locus whose ANS/RNS had the most significant association with age. **(g)** The locus whose ANS/RNS differed most significantly between men and women. The data in the box range from 25% to 75%. Error bars indicate SD×1.5. STC: Superior Temporal Cortex (See Fig. 2 for the full names of the remaining regions).

We calculated the overall (averaged across all voxels) and regional (averaged across the voxels within each part in Supplementary Fig. 4) ANS and RNS of LH and RH for each of the healthy participants (6 individuals had no age and sex information, we only analyzed n = 903 individuals here). Then, we regressed gender and scanner information from the ANS/RNS when analyzing the effect of age and regressed age and scanner information from the ANS/RNS when analyzing the effect of gender. Interestingly, the overall ANS of LH and RH showed a positive correlation with age (p values < 10^-10^, Spearman correlation; Fig. 3b), whereas the overall RNS of LH and RH showed a negative correlation with age (p < 10^-10^, Spearman correlation; Fig. 3b). The overall hemispheric ANS and RNS were higher in males than in females (p values < 0.01, two-sample t test; Fig. 3c). Fig. 3d shows the correlations (r values) between age and each regional ANS/RNS, and Fig. 3e shows the differences (t values) between males and females for each regional ANS/RNS. These line graphs all surprisingly indicated an almost upward trend rather than a random distribution, suggesting that the ANS and RNS may be used to guide the parcellation of the brain.

Moreover, we performed voxelwise analyses to identify loci closely associated with age and sex (Fig. 3f and g). We found that the RNS was more sensitive to sex differences than the ANS, and both the ANS and RNS were sensitive to changes in age. Significant areas were located mainly in the cingulate gyrus, insula, subcortex, and cerebellum.

Together, these results demonstrate that the hemispheric ANS and RNS are meaningful biomarkers that are closely related to the basic human phenotype.

### ANS and RNS are associated with cognition, the environment, and disease

Individual cognitive function is affected by brain lateralization [30, 34], the external environment is an important cause of brain lateralization [35], and alterations in the typical brain lateralization lead to diseases [9, 10, 34–36]. In this section, to further examine whether the ANS and RNS are an accurate representation of hemispheric specialization, we analyzed the association between the ANS and RNS with cognitive measures, demographic information, and brain diseases.

The Cam-CAN database provides behavioral measures of participants, and the CNP database provides intelligence tests of participants. We used a multivariate explanatory model [37] (see Methods) to estimate the association between the ANS and RNS with these metrics and used the actual hemisphere as a baseline. The results indicated that both ANS and RNS of LH and RH can accurately explain the cognitive abilities of individuals (p < 10^-30^, hereditary analysis; Fig. 4a and b). Moreover, except for the ANS of the LH in CNP, the remaining ANS and RNS can explain cognitive abilities better than actual GM (p values < 10^-10^, paired sample t test; Fig. 4a and b).

**Fig. 4.**
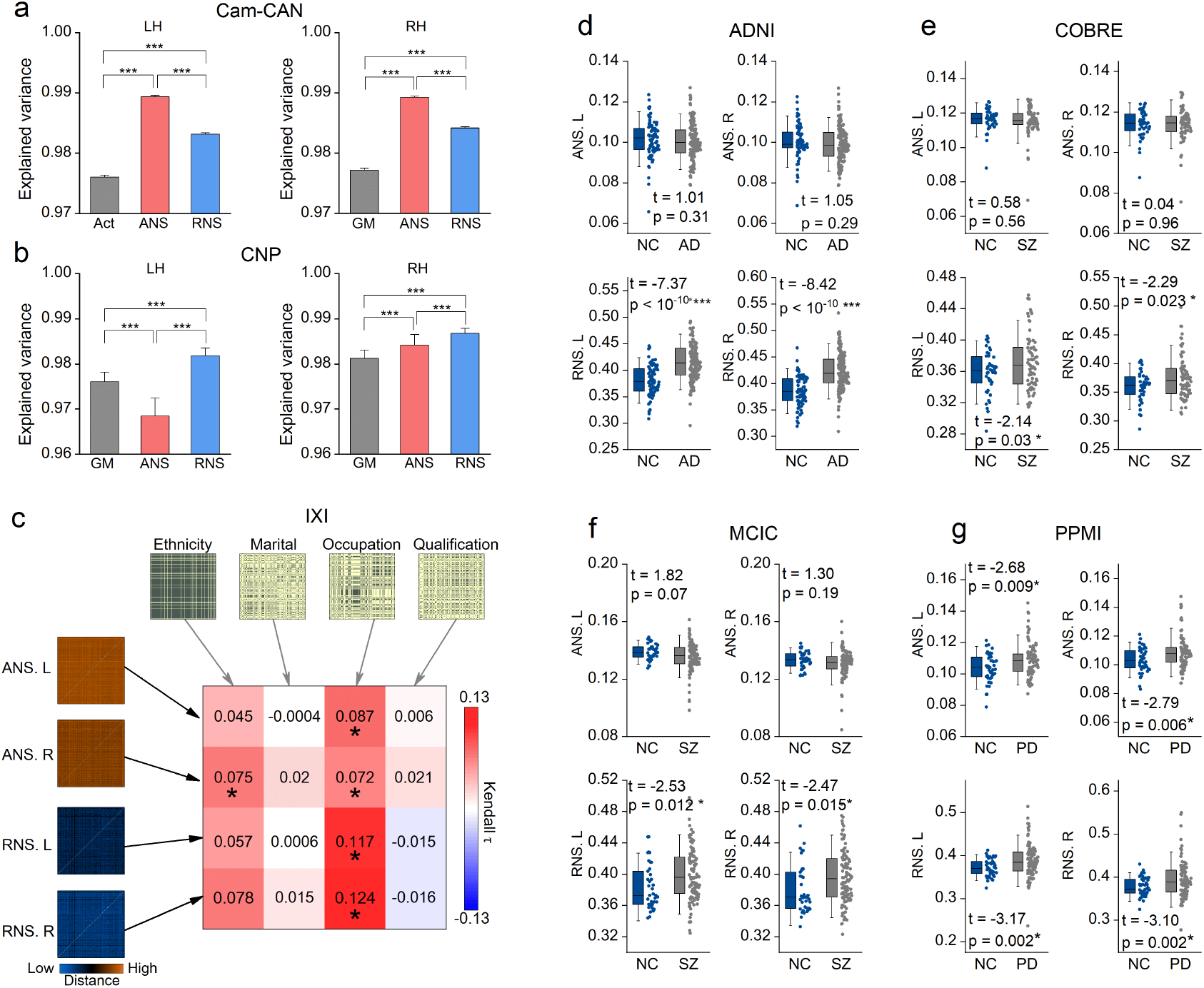
Association between ANS/RNS and cognitive ability, environmental factors and disease. **(a)** and **(b)** Behavioral performance and intelligence measures explained by hemispheric ANS, RNS, and actual hemisphere. The data used in (a) (n = 232) and (b) (n = 50) are collected from the Cam-CAN and CNP datasets, respectively. The explained variance is calculated by a multivariate explanatory model. Paired sample t tests were used for comparisons between groups. The error bar indicates the SD×1.5. *** indicates p < 10^-10^. **(c)** RSA between ANS/RNS and demographic information in the IXI dataset (n = 206). Kendall rank correlation was used to estimate the similarity between RMDs. * indicates p < 0.05 with 5000 permutation tests. **(d) - (g)** Comparison of overall hemispherical ANS/RNS between patients and healthy individuals from the ADNI (154 AD, 82 NC), COBRE (74 SZ, 44 NC), MCIC (104 SZ, 37 NC), and PPMI (98 PD, 45 NC) datasets (two-sided two-sample t test). The data in the box range from 25% to 75%. Error bars indicate SD×1.5. AD: Alzheimer’s disease. SZ: Schizophrenia. PD: Parkinson’s disease.

Next, we explored the relationship between ANS/RNS and demographics (e.g., ethnicity, marriage, occupation, and qualification) of participants in the IXI dataset using RSA [38]. We found that occupation was associated with both ANS and RNS of LH and RH, and ethnicity was associated with ANS of RH (p < 0.05, 5000 permutation test). This result suggests that ANS and RNS are indeed under the influence of some specific environmental factors.

To investigate whether diseases affected the ANS and RNS, we analyzed patient data from four datasets, including Alzheimer’s disease (AD), schizophrenia (SZ), and Parkinson’s disease (PD). We also calculated the overall ANS and RNS of the LH and RH for patients and then compared healthy individuals and patients in each database separately. The results showed that only the overall hemispheric ANS of PD patients was significantly higher than that of healthy subjects (p < 0.05, two-sample t test on PPMI, Fig. 4g), but the overall hemispheric RNS was significantly higher in all patients (p < 0.05, two-sample t test; Fig. 4d-g). In the comparison of regional ANS/RNS, we found more between-group differences (Supplementary Fig. 7). The regional ANS/RNS alterations were not identical for different types of diseases.

These findings further suggest that the ANS and RNS can capture interindividual variation caused by cognitive function and environmental factors and abnormalities in brain structure. In the future, they may be used as biomarkers to infer the characteristics of individuals.

### Spatial autocorrelation of ANS and RNS

Brain features possess spatial autocorrelation (SA) [39]. Due to SA, values of brain features in spatially proximal regions tend to be more similar than values of spatially distant regions. To demonstrate whether the spatial distributions of the ANS and RNS maps are biologically plausible, we explored the SA of the ANS and RNS. We randomly selected approximately 70,000 voxels and calculated the partial correlation (age, gender, and scanner as covariates) between each voxel and its surrounding voxels at distances ranging from 1 to 3. The results showed that for both ANS and RNS, the correlations decreased significantly with increasing distance (p values = 0, one-way ANOVA; Fig. 5a and b), suggesting that the spatial distributions of ANS and RNS maps follow SA. The correlation between the ANS/RNS of voxels and the GM density of voxels around them also decreased with increasing distance (p values < 10^-10^, one-way ANOVA; Supplementary Fig. 8).

**Fig. 5.**
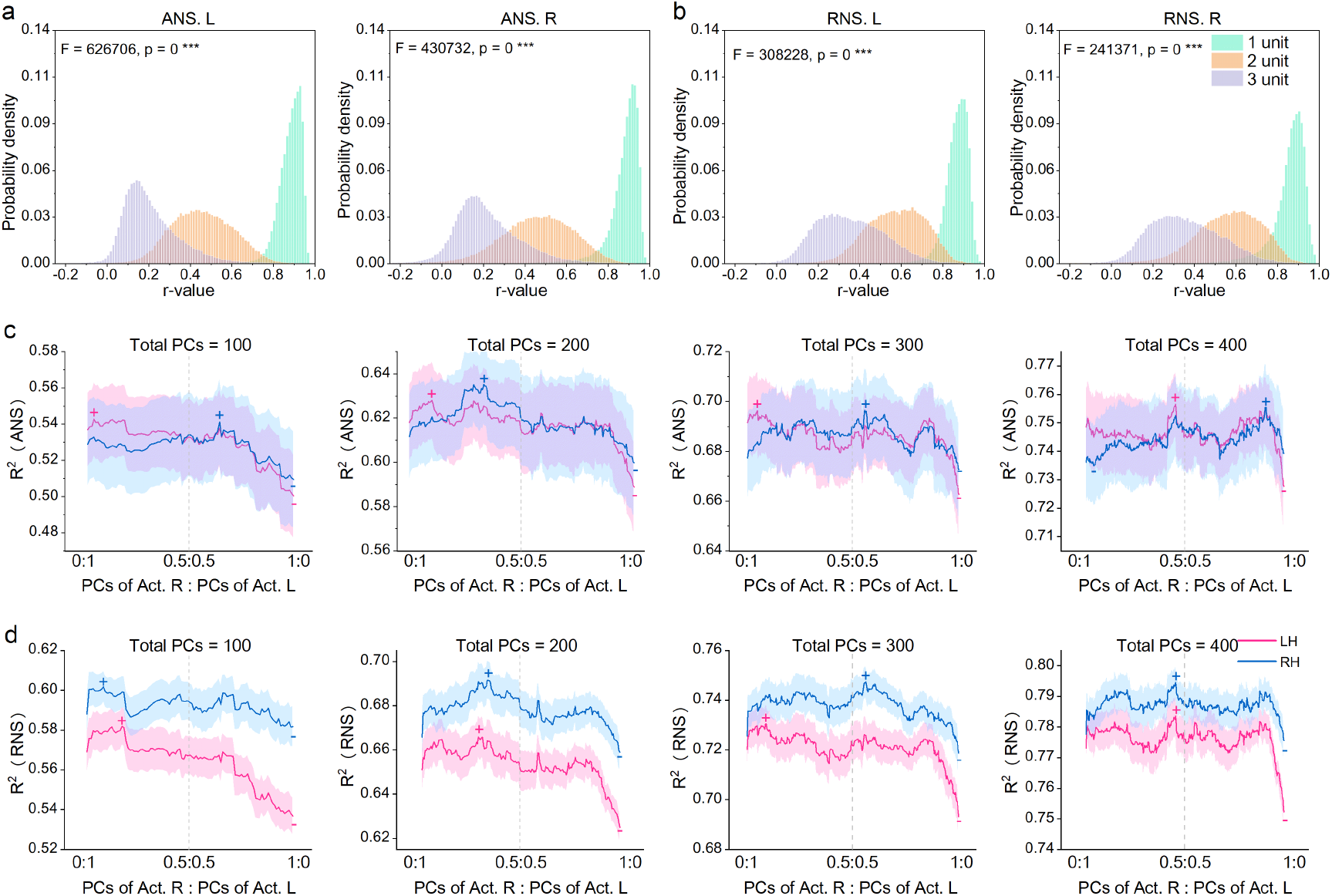
Spatial distribution properties of ANS/RNS and the effect of LH and RH on ANS/RNS. **(a)** and **(b)** Correlation of the ANS/RNS of each voxel (n = 70000) with the ANS/RNS of the voxel at a distance K from it. The unit of distance is the length of a voxel (1.5 mm). In this study, K = 1, 2, and 3. The X-axis represents the r-value, and the Y-axis represents the probability density. One-way analysis of variance (ANOVA) was used for comparisons between groups. **(c)** and **(d)** Regression analysis between the hemispheric actual GM and the hemispheric ANS/RNS. The independent variable is the PCs of actual GM, and the dependent variable is the mean ANS/RNS within each part (Supplementary Fig. 4). The X-axis represents the ratio between PCs of LH and PCs of RH in the independent variable, where the total number of PCs is constant. The Y-axis represents the mean R^2^ across 10 parts, which evaluates how well the model fits. The shaded part indicates the standard error (SE). (+) indicates the maximum value. (–) indicates the minimum value.

### Association of brain structural patterns with hemispheric specialization

We have demonstrated the reliability and plausibility of the hemispheric ANS and RNS in several ways. Here, we analyzed the association between brain structural patterns and hemispheric specialization. Multiple linear regression was used to map the representations of actual GM to the hemispheric ANS/RNS. As shown in Fig. 2a, we first extracted the PCs of the actual LH and RH (the transformation matrix was estimated using the training set, which was later applied to the validation set). Then, keeping the total number of PCs as N (in this study, N = 100, 200, 300 or 400), we concatenated the PCs of LH with those of RH in different proportions (ranging from 0:1 to 1:0 with a step of 1/N) and then used each combination to fit the ANS/RNS for each of the 10 parts (Supplementary Fig. 4). Fig. 5c and d show the averaged R^2^ of fitting of each combination. The models fitted the ANS of RH and LH essentially identically (the red and blue lines overlap to a large extent, Fig. 5c) but fitted the RNS of the RH better than that of the LH (the blue line is higher than the red line, Fig. 5d). Additionally, the fit of the model decreased significantly when the ratio between PCs of RH and PCs of LH was close to 0:1 (the smallest R^2^ values were all located on the rightmost side of the X-axis, except for the ANS of LH at total PCs = 400). These findings suggest that the formation of RH specialization is more likely to be influenced by innate brain patterns and that more biological factors contributing to hemispheric specialization are in the LH.

### Important areas for controlling hemispheric specialization

In this section, we performed voxelwise analysis to identify loci that were closely related to hemispheric specialization. As we did for Fig. 2c and d, we computed the first two PCs of ANS/RNS maps of LH and RH across healthy participants (n = 909). Next, we calculated the correlations between two PCs with the actual GM density of each voxel (Pearson partial correlation using age, sex, and scanner as covariates). Fig. 6a and b show the loci most significantly correlated with the ANS and RNS, respectively. The results suggest that the cingulate cortex, insula, parahippocampal gyrus, cerebellum, and subcortical regions (hippocampus and putamen) are the most important areas for controlling hemispheric specialization. These significant regions largely overlapped with the major variant loci in the ANS and RNS (Fig. 2c and d, and Fig. 3e and f).

**Fig. 6.**
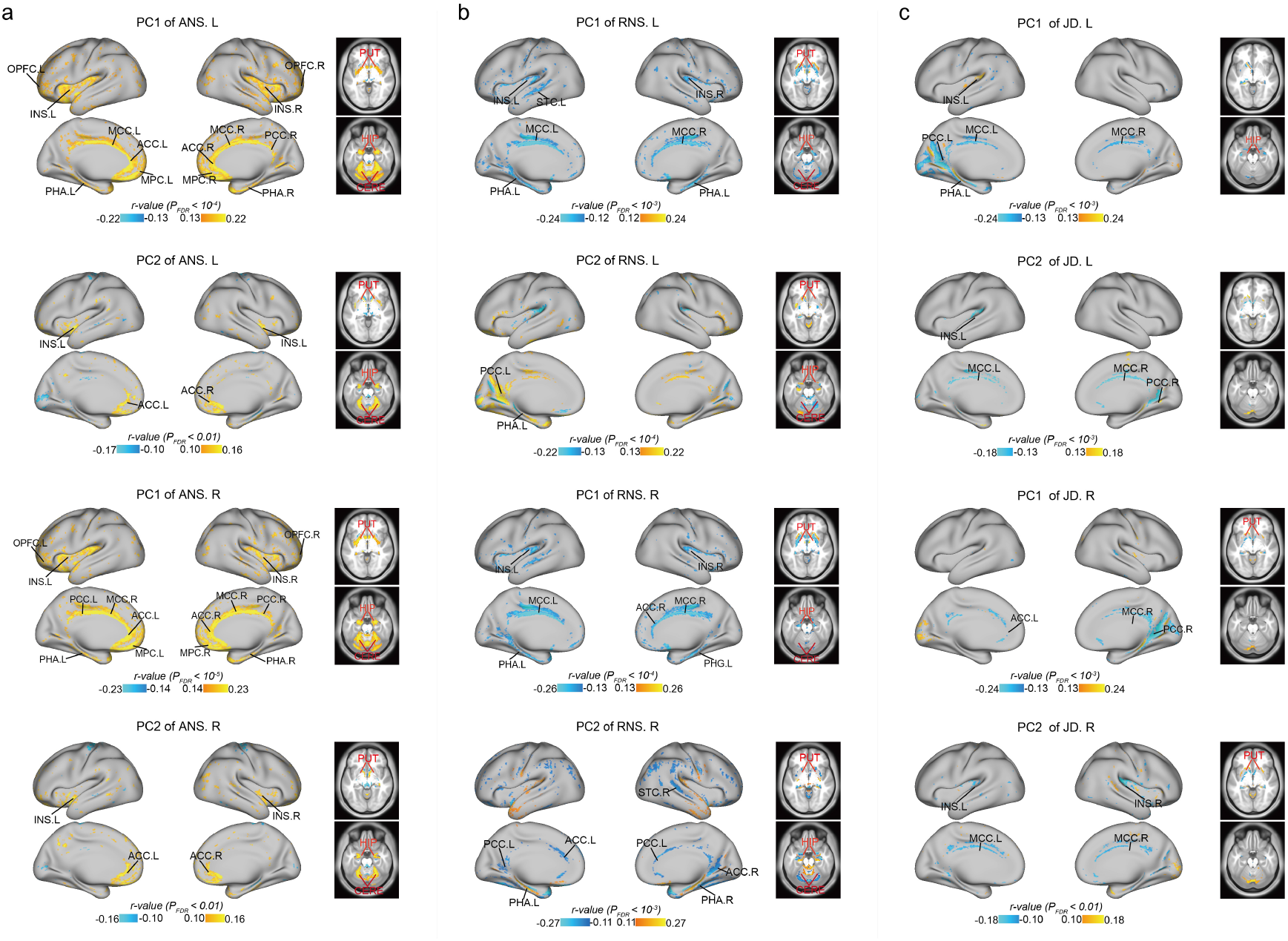
Visualization of important regions that control and drive hemispheric specialization. **(a)** and **(b)** Correlation between the first two PCs of the hemispherical ANS/RNS maps with each actual voxel. **(c)** Correlation between the first two PCs of the hemispherical Jacobian determinant (JD) maps with each actual voxel. Pearson partial correlation analysis was used (age, sex, and scanner were used as covariates). The p values were corrected by FDR. In different subgraphs, we used different thresholds to locate the most significant voxels. OPFC: Orbital and Polar Frontal Cortex, MPC: Medial Prefrontal Cortex (See Fig. 2 and 3 for the full names of the remaining regions).

Furthermore, we identified the loci closely associated with the process of hemispheric specialization. First, we calculated the deformation field using a nonlinear registration that transformed the reconstructed hemispheres into the corresponding actual hemispheres. We then computed the Jacobi determinant (JD) of the deformation field as an index of local expansion or contraction (Supplementary Fig. 9) [40]. The resulting JD map can be used to estimate the dynamics leading to hemispheric specialization formation (How to drive homologous components to form an actual hemisphere). Consistent with the above analysis, we computed the first two PCs of the LH and RH JD maps and calculated the correlation between PCs with the GM density of each voxel. The significant areas (Fig. 6c) were largely consistent with those in Fig. 6a and b suggesting that they play a dominant role in driving hemispheric specialization. We also examined whether aging, sex, and environmental factors alter hemispheric specialization dynamics. We found that only PC2 of the JD map of RH was significantly correlated with age (r = −0.195, p = 3.2×10^-9^, Spearman correlation; Supplementary Fig. 10), and there was no difference between males and females (p>0.05, two sample t test, Supplementary Fig. 10). RSA (similar to Fig. 4b) showed that JD maps were not associated with demographic information (p > 0.05, 1000 permutation test). Thus, this result demonstrates a relatively stable direction of individual hemispheric specialization, which is only slightly influenced by aging.

## Discussion

Our study proposed a new method using a self-supervised DGN to reveal hemispheric specializations in the brain. The results suggest that our method is stable, reliable, and plausible and is able to capture individual variation characterizing a wide range of individual traits. We expect that our method will provide inspiration for the characterization of brain function and structure. Next, we discuss our findings from the joint perspective of methodology, neuroscience, and applications.

Deep learning provides powerful and promising tools for neuroimaging research, as it can learn high-level and abstract representations of data using complex nonlinear transformations [41]. Deep learning methods have been widely used for medical image classification, segmentation, alignment, and other tasks [42]. Moreover, recently researchers have focused more on using deep learning methods to automatically capture and identify high-level biomarkers. Two related studies demonstrate that the latent variables and residuals generated by variational autoencoders (VAE) accurately captured the interindividual variation in the fMRI data [43, 44]. Chavas et al. suggested that β-VAE can help reveal unknown cortical folding patterns [45]. Zhao et al. proposed a new autoencoder model to obtain representations of brain function [46]. Aglinskas et al. leveraged contrastive variational autoencoders (CVAEs) to better characterize the neuroanatomical variation specific to autism spectrum disorder (ASD) [47]. These studies and our work share the same inspiration: using self-supervised deep learning methods to obtain biologically meaningful markers. However, instead of encoding the entire brain image with an autoencoder-based model, our work uses masked image modeling (MIM) to create a mapping between the two hemispheres to reveal biologically specific components of each part. It has been suggested that the learned representation in autoencoders is likely to be the compression of image content, while MIMs have a deeper semantic understanding of the scene[18, 48]. Biomedical data collection is time-consuming and laborious, so acquiring training targets for complex tasks is often impossible, but the results demonstrate that the MIMs are able to learn the information we expect without target samples. This study supports and extends previous theory to some extent, showing that MIM approaches are promising for deep characterization of the brain. Another difference is that most autoencoder-based representation learning methods use the latent space as a new biomarker, which is informative but hard to interpret. However, in this work, the representations restored in the original image space are still informative and easy to understand, which facilitates subsequent analysis at the region or voxel level.

In addition, we note that our workflow is theoretically similar to that of the activity flow mapping method [49]. Activity flow mapping shows that task-evoked activity in one region can be predicted by the FC-weighted sum of task-evoked activity in other regions and the similarity between predicted and actual activity can be used to estimate interregional information transfer and reveal disease-related features [50, 51]. Similarly, our study essentially uses one region to explain another region, after which the unexplained variance is considered to be a biological indicator. Neurons in the brain are all interdependent [52], which provides a biological possibility for uncovering the mapping between brain regions. Our work and previous studies demonstrate that when the established mapping function is biologically plausible, its derivatives are likely to reveal high-level biological features. In terms of performance, however, deep learning methods have greater potential to fit intricate interactions between neurons than simple linear methods.

To estimate voxelwise or vertexwise lateralization, most studies have used significant warping and spatial smoothing to force the brain into a symmetric space, thereby establishing a left-right correspondence. Then, the lateralization is quantified by comparing the original image and the flipped image. However, even if the brain is spatially symmetric, this does not guarantee a biological correspondence because the global pattern of the brain is continuously reshaped under the influence of external factors [7]. In addition, a fair comparison usually requires the condition that the LH and RH are within the same data space. However, the structure and function of the brain already exhibit significant asymmetries at birth [53, 54]. This implies that the neurobiological substrates of the LH and RH (similar to the developmental functions *f1* and *f2* in Fig. 1a) are already heterogeneous at the beginning of life, some of which are unexplainable, confounding many attempts at alignment.

In contrast, in our work, the generator reveals the complex dependencies between homologous components of the two hemispheres, and the discriminator facilitates learning the distribution of target spaces conditioned on source spaces. Our approach implements semantic transfer, which expresses the content of one hemisphere in the style of the opposite hemisphere. This process eliminates gaps between different spaces, and establishes a biological correspondence between LH and RH, thus allowing for a fair comparison.

In this study, we proposed two metrics for estimating hemispheric specialization: ANS and RNS. Our results demonstrate that they both encode much biological information, such as age, gender, disease, environmental factors, and cognitive function. However, through a comprehensive comparison, we believe that the RNS is more reliable and informative. The RNS is calculated from the ratio of the ANS to GM density, and this operation mitigates the interference caused by some confounding factors. For example, the ANS was negatively correlated with age while the RNS was positively correlated with age. This is because the RNS eliminates the effect of GM density decreasing with age. Furthermore, this finding suggests that the proportion of specific components underlies brain asymmetry and dominates the formation of interindividual differences.

Evidence has shown that the cingulate gyrus, insula, subcortex, and cerebellum have strong anatomical and functional connections to brain structures and possess important hub-like properties [55–57]. In the current study, these areas had relatively low ANS and RNS, implying their robust structure and biological symmetry, but their ANS and RNS accounted for most of the variation between individuals and correlated strongly with sex and age. Furthermore, these regions controlled the overall pattern of hemispheric specialization. Our results support and extend previous findings suggesting that these areas play an important role in the establishment of hemispheric specialization and, to some extent, reveal the origin of hemispheric specialization.

In this work, our main goal is to suggest that DGNs are promising tools for the study of brain asymmetry. We used only a simple network backbone consisting of several CNN blocks for validation. In the future, researchers can modify the network backbone and the loss function according to the needs of the study, further improving the accuracy of the results. Examples include using a hybrid framework of CNN and self-attention [58] to focus on both local and global information or adding a regularization module to capture more semantic information. In addition, the results provided a preliminary indication that our proposed ANS and RNS capture and encode a variety of biologically meaningful information. They can be extended and applied to a wide variety of research as complementary tools of neuroscience, such as for revealing the properties of certain regions of interest (ROIs), exploring the development and aging of the brain, and diagnosing brain diseases. Moreover, in this study, we mainly performed a global analysis that only considered the absolute value of the ANS or RNS. However, the positive and negative nature of the model error also has important biological implications, which have been demonstrated in brain age-related studies [59]. We will later clarify this issue to further improve the interpretability of our method.

Although our method is oriented toward brain neuroanatomy, it can be easily transferred to fMRI analysis by simply altering the architecture of the network. The analytical process in our work is not limited to the study of hemispheric specialization but can be used to study the specialization of any region, as long as there is a sound neuroscientific basis for it. For example, by establishing a mapping between cortical and subcortical tissue, we can infer their specialization components and processes.

In conclusion, we leveraged a self-supervised DGN to disentangle the features of hemispheric specialization. The results demonstrate that this approach is feasible and promising. Our study not only explores brain asymmetry from a new perspective but also sheds new light on the application of DGN to neuroimaging.

## Methods

### MRI data

All neuroimaging data used in the study were T1-weighted MRI scans. We collected data on healthy people and patients from 14 publicly available datasets, made available via various data-sharing initiatives; there were a total of 2852 samples, with an age range of 4-90.4 years (Table.1). Among the datasets, ten datasets were used to train the models and analyze the main results. One dataset was used to evaluate the generality of the deep learning model. Three datasets were used to evaluate within-scanner reliability, between-scanner reliability, and heritability. The imaging parameters are shown in Supplementary Table 3.

The datasets used to train the model and analyze the main results include Beijing Zang, IXI, Cam-CAN [60], COBRE, MCIC [61], ADNI [62], NKI, NUSDAST [63], CNP [64], PPMI [65]. According to the age and gender distribution and sample size of each dataset, 1372 healthy subjects were randomly selected to train the models. The remaining 909 healthy subjects and 430 patients were used for the analysis of the main results.

The ICBM with 78 healthy subjects was used to evaluate the convergence of the model and the quality of the reconstructed images during training. When the loss value and the quality of the reconstructed image stabilized, we stopped the training and saved the model.

The MCS [66] and Yale-TRT [67] datasets were used to evaluate the stability of the same individual in different scans. Each health participant in MCS underwent 2 scanning sessions on separate days, which were collected using a same 3T Siemens Trio. Each health participant in Yale-TRT underwent scans over 4 sessions approximately 1 week apart, using the 2 identically configured Siemens 3 T Tim Trio scanners: 2 sessions used “Scanner A”, and the other 2 sessions used “Scanner B”. We used two scans of 10 subjects in MCS and one scan from “Scanner A” and one scan from “Scanner B” of 8 subjects in Yale-TRT.

The QTIM [68] was used to evaluate the heritability among family members. It is a multimodal neuroimaging dataset of young adult twins and siblings. In our study, we randomly selected 47 subjects from 22 families.

### Data preprocessing

We performed minimal preprocessing on the data. The T1-weighted images were segmented into separate tissue classes. All gray matter (GM) density maps were normalized to the standard Montreal Neurological Institute (MNI) template with a 1.5 × 1.5 × 1.5 mm^3^ voxel resolution. These steps were implemented in FSL. We then used a GM probability mask (GM probability threshold = 0.15) to extract voxels corresponding to GM tissue, removing the noise and artifacts at the edge of the GM tissue. Finally, we split the GM into left and right hemispheres along the X = 0 axis in the transverse section and cropped each hemisphere to remove the black background.

Data were excluded if an anatomical scan failed to complete the tissue segmentation procedure or if tissue segmentation was inaccurate. The number of subjects stated above was determined after quality assessment (Table. 1).

### Deep learning model

The GAN training strategy is to define a game between two competing networks. The generator network (G) transforms data from one space to another. The discriminator network (D) receives either a generated sample or a real data sample and must distinguish between the two. G is trained to produce more realistic images, eventually confusing D. The context encoder (CE) is a self-supervised masking image model (MIM) based on the GAN architecture[18]. Its G consists of an encoder, which captures the context of an image into a compact latent feature representation, and a decoder, which uses this representation to generate the missing image content. D allows G to capture more details and semantic information, improving the quality of the image reconstruction.

In this study, the encoder consists of four repeated down-block of a 3D convolution neural network (CNN) layer, a 3D batch-normalization (BN) layer, and a rectified linear unit (ReLU). The decoder consists of four repeated up-block of a 3D deconvolution layer, a 3D BN layer, and a ReLU. A channel wise fully-connected layer is used to connect the encoder and the decoder, allowing each unit in the decoder to analyze about the entire image content. D consists of four CNN layers and one classification layer.

The LH and RH are described by the following formulations:

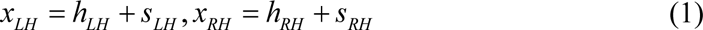

where *h* indicates homologous components, *s* indicates specific components. Both *h_LH_* and *h_RH_* originate from shared factors, thus they are not independent:

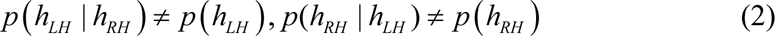

However, *s_LH_* and *s_RH_* are derived from factors unique to the LH and RH, respectively, so they are independent:

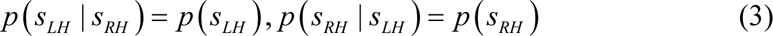

As a result, the network can still estimate the *h* in one hemisphere by understanding the opposite hemisphere.

The objective of the network is to use the LH or RH to predict the RH or LH, respectively, thus naturally revealing the innate mapping between *h_LH_* and *h_RH_*. The loss function is a combination of reconstruction loss and adversarial loss. The reconstruction loss is responsible for capturing the overall structure of the missing region and coherence regarding its context but tends to average together the multiple modes in predictions. We used a voxelwise L2 distance as the reconstruction loss. Take using the LH to predict the RH as an example:

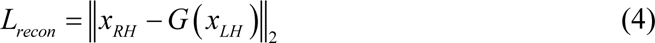

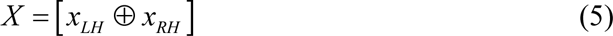

where X is the whole brain. ⊕ indicates the concatenation operation.

The adversarial loss, on the other hand, tries to make the prediction look real and has the effect of picking a particular mode from the distribution[19]. Inspired by the Wasserstein GAN (WGAN), we add a gradient penalty to our adversarial loss, which can improve the stability of training, eliminate problems such as mode collapse, and provide meaningful learning curves useful for debugging and hyperparameter searches [69]. The adversarial loss for context encoders is:

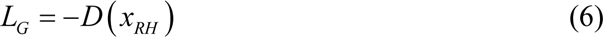

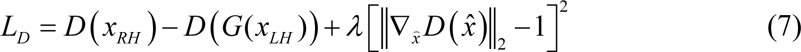

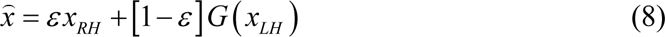

where ∇ stands for gradient calculation. *ε* is a random number between 0 and 1. We define the overall loss function as:

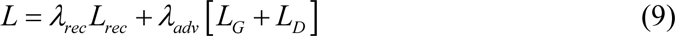

In this work, *λ_rec_* is 0.9 and *λ_adv_* is 0.1. We trained two CE models by the same strategy, one using the LH to predict the RH, and the other using the RH to predict the LH. The models were trained on four NVIDIA Tesla P100 GPUs. The training epoch was 3000 when the reconstruction reached stability. We evaluated the quality of the reconstructed images based on a combination of loss values and subjective observations.

### The measure of hemispheric specificity

We fed the LH (resp. RH) image into the trained model and obtained the reconstructed RH (resp. LH) image. The hemispherical ANS map is represented by a voxelwise distance between the actual and the reconstructed image.

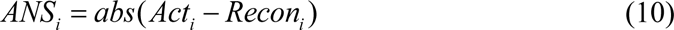

The index *i* indicates LH or RH. In the training process, the objective is to reduce the distance between the outputs and the objects. Therefore, we do not consider the positive or negative of the difference between the reconstructed voxel and the voxel.

Furthermore, we calculated the relative neuroanatomical specificity (RNS) map, estimating the proportion of specific components in the hemisphere.

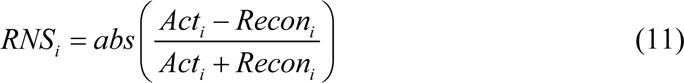

To explore whether the biological information described by the different ranges of ANS/RNS is varied, we divided the voxels into different parts. According to the average ANS or RNS map across individuals, we ranked voxels in descending order and divided them into ten equal parts. For example, the first part corresponds to the voxels with the top 10% ANS or RNS, and the rest can be understood in the same manner.

### Verification of robustness, generalization, and heritability

To verify that the reconstructed images capture the intrinsic characteristics of each individual, and not just a rough description of the brain shape of the group, we calculated the similarity between each reconstructed image and the corresponding actual image, as well as the noncorresponding actual images. The image was first reshaped into a one-dimensional vector. Pearson correlation analysis was used to calculate the similarity between vectors. Two-sample t tests were used to compare the r-values between corresponding images and those between noncorresponding images.

We used the ‘virtual lesion’ method to verify that our model accounts for the characteristics of all brain regions, rather than only using some regions to establish mapping. The AAL atlas [26] was used to define the regions of interest (ROIs). We masked one ROI (assigned a value of 0 to the voxels within the region) and then inputted the damaged image into the trained model to obtain the reconstructed image. Each ROI performed the above step. Furthermore, we compared the reconstruction losses from damaged images with those from complete images.

To verify that the ANS and RNS are reliable indicators, we analyzed the stability of the same individual in different scans collected by the same or different scanners. We used Pearson correlation analysis to calculate the similarity between subjects and within subjects across scan sessions.

To verify that ANS and RNS can be inherited, we analyzed the consistency among family members, including twins and siblings. The Pearson correlation coefficient was used to estimate the similarity within and between families.

### Principal component analysis

Principal component analysis (PCA) was used to reduce the dimensionality of features and to indicate major interindividual variation. For the actual hemispheres (GM maps), the transformation matrices of the PCA were estimated by the individuals in the training set (n =1372) and then were applied to individuals in the validation set. For the ANS, RNS, and Jacobian determinant maps, the transformation matrix obtained by 99% of the subjects in the validation set was used to calculate PCs for the remaining 1% of subjects. PCA was performed separately on the LH and RH.

To identify loci of individual variability related to anatomical specificity, we calculated the first two principal components (PCs) that account for the most variation in the ANS, RNS, or Jacobian determinant maps. To identify areas associated with each PC, we correlated the PC loadings to the ANS, RNS, or Jacobian determinant values for each voxel.

### Regression analysis

Regression models were used to map actual hemispheres to age or ANS and RNS. When predicting ages, PCA was used to reduce the dimensions of the GM maps, for both the LH and RH. We then concatenated the PCs of the LH and RH, explaining 50% of the variation, and used a linear regression model to establish the mapping between PCs and age. These steps were performed on the individuals in the training set (n=1372). After training the models, the transformation matrix of the PCA and the weights in the linear regression model were used to predict the age of healthy individuals in the validation set (n=909). To explore whether the reconstructed images can properly depict brain aging, we applied four sets of images, (1) *Act_LH_* ⊕ *Act_RH_*, (2) *Act_LH_* ⊕ *Recon_RH_*, (3) *Recon_LH_* ⊕ *Act_RH_* and (4) *Recon_LH_* ⊕ *Recon_RH_*, to predict ages. *Act* denotes the real image. *Recon* denotes the generated image. The subscript denotes the index of the hemisphere. ⊕ indicates the concatenation operation.

When predicting ANS and RNS, we also applied the transformation matrix from the training set to reduce the dimension of the actual GM within the validation set. Then, we mapped the PCs of actual GMs to ANS or RNS using multiple linear regression models, and measured fittings according to R^2^. Specifically, age, sex, and scanner information were regressed out from PCs and ANS or RNS. We concatenated the PCs of the actual GM of LH and RH in different proportions, ranging from 0:1 to 1:0 while keeping the total number N of PCs unchanged. For example, a ratio of 0.5:0.5 indicates concatenating the top N/2 PCs of the LH and top N/2 PCs of the RH; a ratio of 0:1 indicates choosing the top N PCs of the RH and no PCs of the LH. Afterward, each combination of PCs was fitted to the mean ANS/RNS of each of the ten parts (Supplementary Fig. 4).

### Representational similarity analysis

We used RSA [38] to examine the associations between neuroanatomical features and the associations between neuroanatomical features and demographic information. We collected the demographic variables from the IXI database, which described the ethnicity, occupation, marriage, and qualification of the subjects. The RDMs of neuroanatomical features was calculated by correlation distance (1 minus the correlation coefficient). For demographic variables, the subject distance was 0 if categorical variables matched (e.g. same ethnicity) and 1 otherwise. Considering that the demographic variables are discrete, we used Kendall rank correlations to estimate the similarity between the RDMs of the demographic variables and the RDMs of the neuroanatomical features. Spearman rank correlation was used to estimate the similarity between the RDMs of neuroanatomical features.

### Variance component model

We used a multivariate variance component model [37] to link anatomical features and behavioral measures. The behavioral measures were collected from the Cam-CAN and CNP datasets. In Cam-CAN, participants completed a battery of behavioral tasks to assess their cognitive functions in the aspects of executive function, emotional processing, motor function, and memory [60]. Our study focused on five tasks including fluid intelligence test, face recognition, picture priming, proverb comprehension, and emotion expression recognition. In CNP datasets, participants took part in the Wechsler Adult Intelligence Scale III (WISN-III) test, which included letter-number sequencing (LNS), matrix reasoning (MR), and vocabulary test (VOC) [64].

We first regressed age and gender from these phenotypic measures, which were then quantile normalized. The multivariate variance component model is shown as follow:

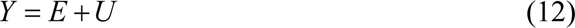

where *Y*, *E*, or *U* is the matrix with a size of *S* × *H*. *S* is the number of individuals, and *H* is the number of behavioral measures (*H* = 5 for Cam-CAN and *H* = 3 for CNP). *Y* contains the *H* processed behavioral measures for all *S* individuals. *E* and *U* represent common environmental factors and unique environmental factors, respectively. *vec*(*E*) ∼ *N* (0, ∑*_e_* ⊗*F*) and *vec*(*U*) ∼ *N* (0, ∑*_e_* ⊗*I*). *F* is a similarity matrix, where *F*(i, j) encodes the anatomical features (GM, ANS, or RNS) similarity of between subjects *i* and *j*. *vec*(.) is the matrix vectorization operator. ⊗ is the Kronecker product of matrices. *I* is an identity matrix, *∑*_*e*_∈ *R*^*H*×*H*^ is a common environmental covariance matrix, and *∑*_*u*_ ∈ *R*^*H*×*H*^is the unique environmental covariance matrix, which is to be estimated from *F* and *Y.* The variance explained by the anatomical features, denoted by *M*, is computed as:

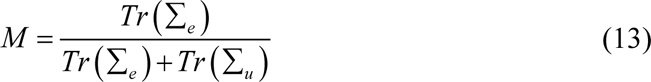

where *Tr*(.) is the trace operator. *M* measures how much inter-individual behavioral variability is explained by inter-individual anatomical features variability.

To facilitate statistical difference between the variance explained by two different metrics, we repeated the above steps *S* times, removing one sample in each calculation. The result *M_-i_* of each time represents the variance explained from whole datasets without subject *i*. Then, a set of *M* values were used for comparison between groups.

### Spatial autocorrelation

To explore whether the spatial distribution of the ANS/RNS was spatially autocorrelated, we analyzed the similarity between voxels at different distances. We first sampled approximately 70,000 voxels at random. We then used Pearson partial correlation to calculate the similarity of a voxel with neighboring voxels at distances from 1 to 3 units (a basic unit is 1.5 mm in this study). We chose the front, back, top, bottom, left, and right voxels of a voxel as its neighboring voxels and averaged them before correlation analysis. One-way analysis of variance was used to compare neighboring voxels at different distances.

### Deformation-based morphometry

We performed deformation-based morphometry analysis to estimate the internal dynamics of hemispheric specialization. Specifically, we applied a nonlinear warping (using inverse-consistent diffeomorphic image registration implemented in ANTs) to transform each reconstructed LH or RH into its corresponding actual hemisphere. The resulting deformation field described how much each voxel was moved from the reconstructed hemisphere to match the corresponding original hemisphere. We then calculated the Jacobian determinant of the deformation field as an index of local expansion or contraction [40]. This estimate captured the local expansion or atrophy required to morph the reconstructed hemisphere to the corresponding actual hemisphere.

### Statistical analysis

Two-tailed paired sample T-test was used to compare different indicators in a set of data. Two-tailed two-sample T-test was used to compare different datasets. One-way ANOVA was used for comparison between multiple groups. Spearman’s rank correlation was used to calculate the association between age and anatomical features. Multiple comparisons were corrected by FDR. Steiger’s z test was used to compare the correlation coefficients (r-values) [70].

## Data and code availability

The ADNI and PPMI datasets are publicly available from https://ida.loni.usc.edu/login.jsp. The COBRE, MCIC, and NUSDAST datasets are publicly available from http://www.schizconnect.org/. The Beijing Zang, ICBM, NKI, and Yale-TRT are publicly available from http://fcon_1000.projects.nitrc.org/. The IXI dataset is publicly available from http://brain-development.org/ixi-dataset/. The Cam-CAN dataset is publicly available from http://www.cam-can.com/. The CNP dataset is publicly available from https://openneuro.org/datasets/ds000030/. The MSC is publicly available at https://openneuro.org/datasets/ds000224/. The QTIM is publicly available from https://openneuro.org/datasets/ds004169/.

All analysis code is available from: https://github.com/book-book24/hemispheric-specializations.git.

## Acknowledgements

This work was supported part by National Natural Science Foundation of China (U20A20191); the Beijing Municipal Science & Technology Commission (Z201100007720009); the Fundamental Research Funds for the Central Universities (2021CX11011); and the China Postdoctoral Science Foundation (2020TQ0040).

## Author contributions

G.W. designed the study, performed the data analysis, and wrote the paper. N.J. built the deep learning framework. Y.M preprocessed the data. T.Y. managed the project and reviewed the paper.

## Supplementary information

**Supplementary Fig. 1.**
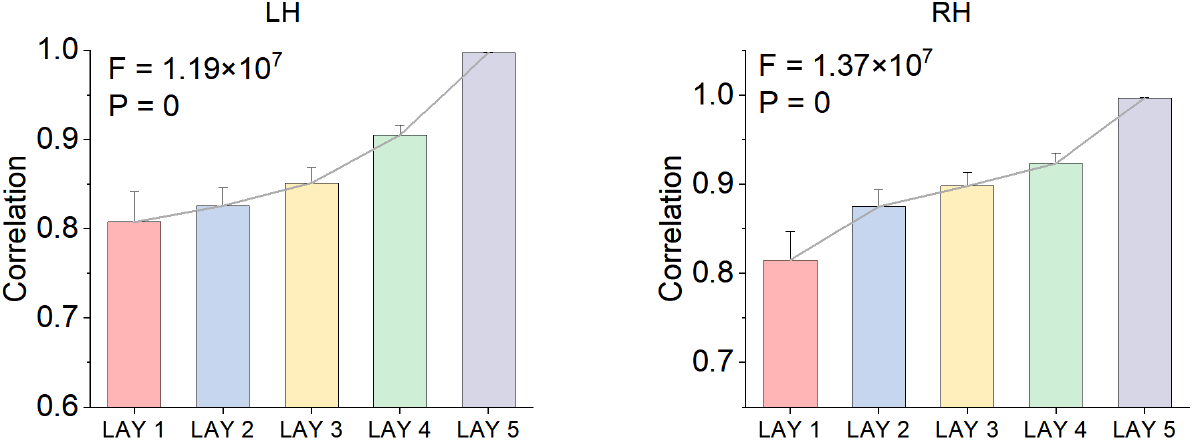
Inter-individual similarity calculated from the feature maps in the models. Data from healthy individuals in the validation set (n = 909). ‘LAY’ denotes the CNN layer in CE. LAY 1 to 4 denote the feature maps extracted from each of the four layers in the encoder, and LAY5 denotes the feature maps extracted from the bottleneck. Error bars indicate SD×1.5.

**Supplementary Fig. 2.**
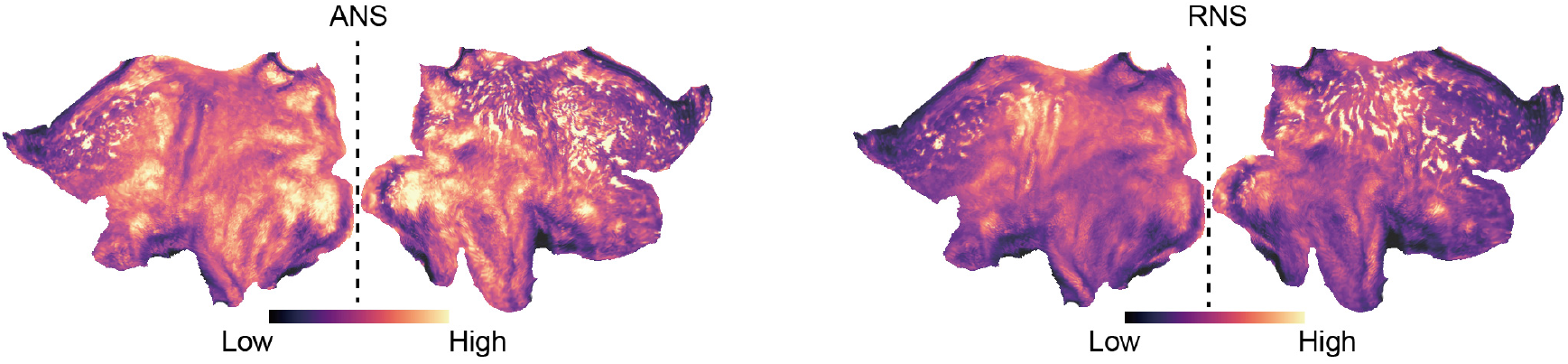
Average hemispheric ANS and RNS visualization in surface space.

**Supplementary Fig. 3.**
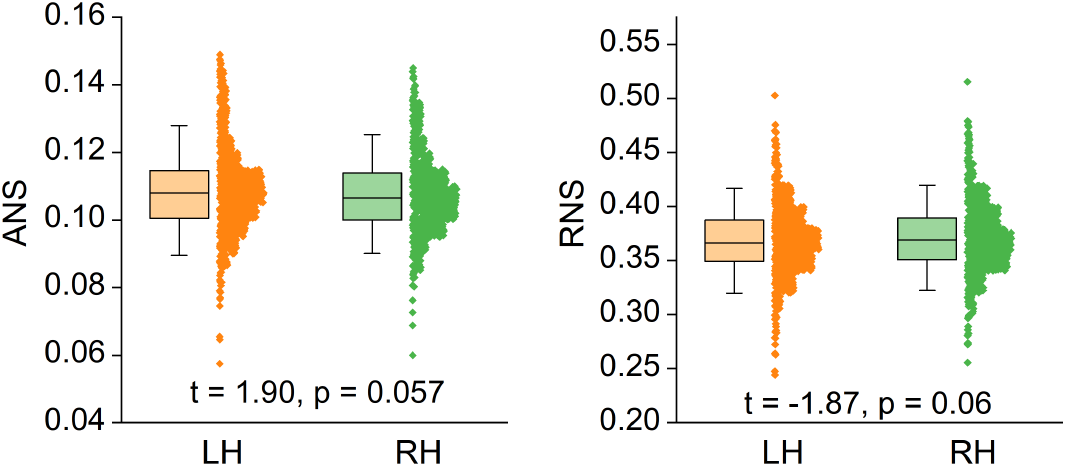
Comparison between the overall ANS/RNS of the left and right brain. Paired samples t-test for comparison between groups. Error bars indicate SD×1.5.

**Supplementary Fig. 4.**
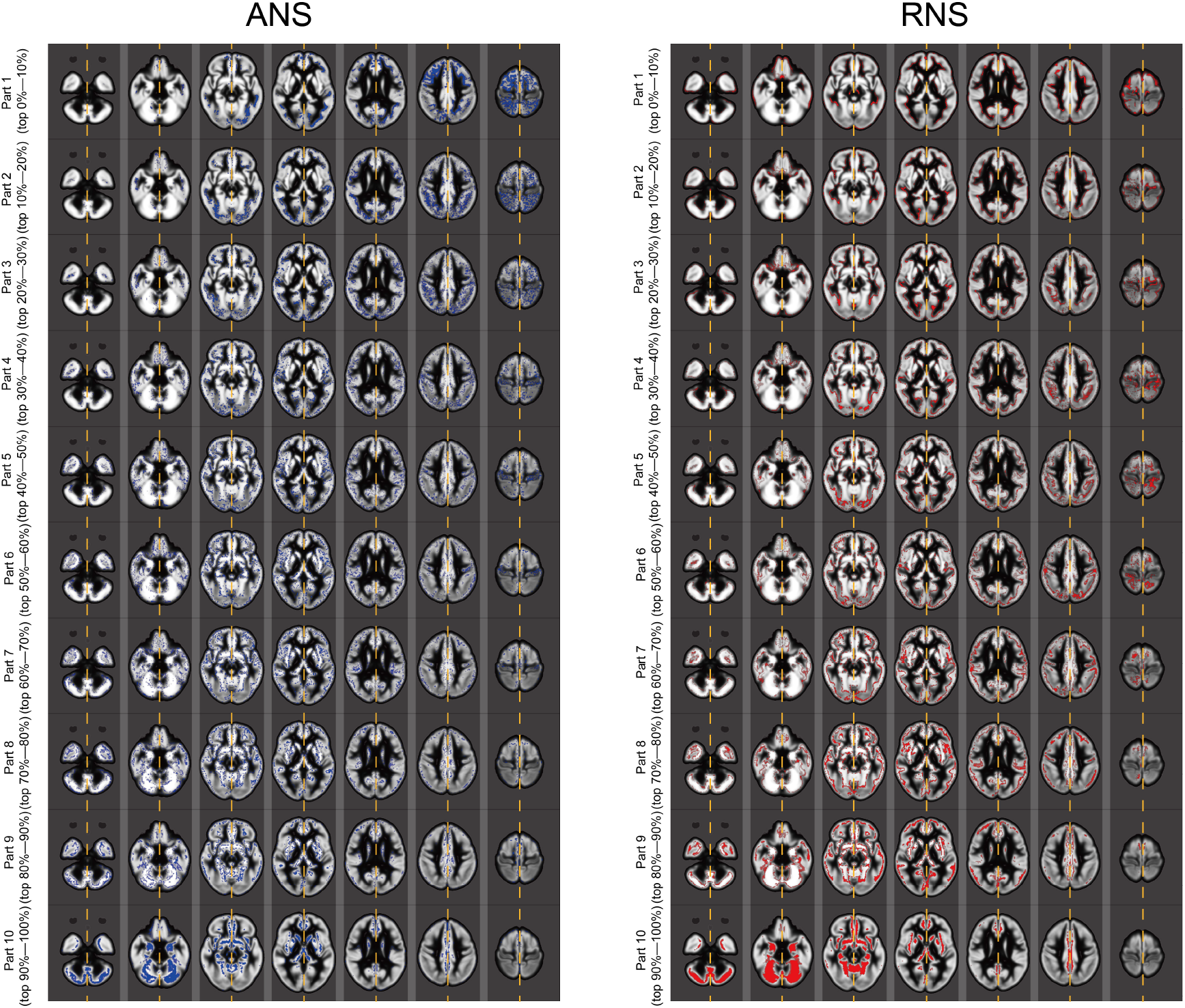
Voxel grouping based on ANS or RNS. According to the value of ANS or RNS, we sort the voxels and divide them into 10 equal parts(regions). For example, region 1 includes voxels in the top 10% of the ANS/RNS ranking, and region 10 includes voxels in the bottom 10% of the ANS/RNS ranking. Of note, the grouping of voxels is performed separately in the LH and RH, but we put the LH and RH together to visualize.

**Supplementary Fig. 5.**
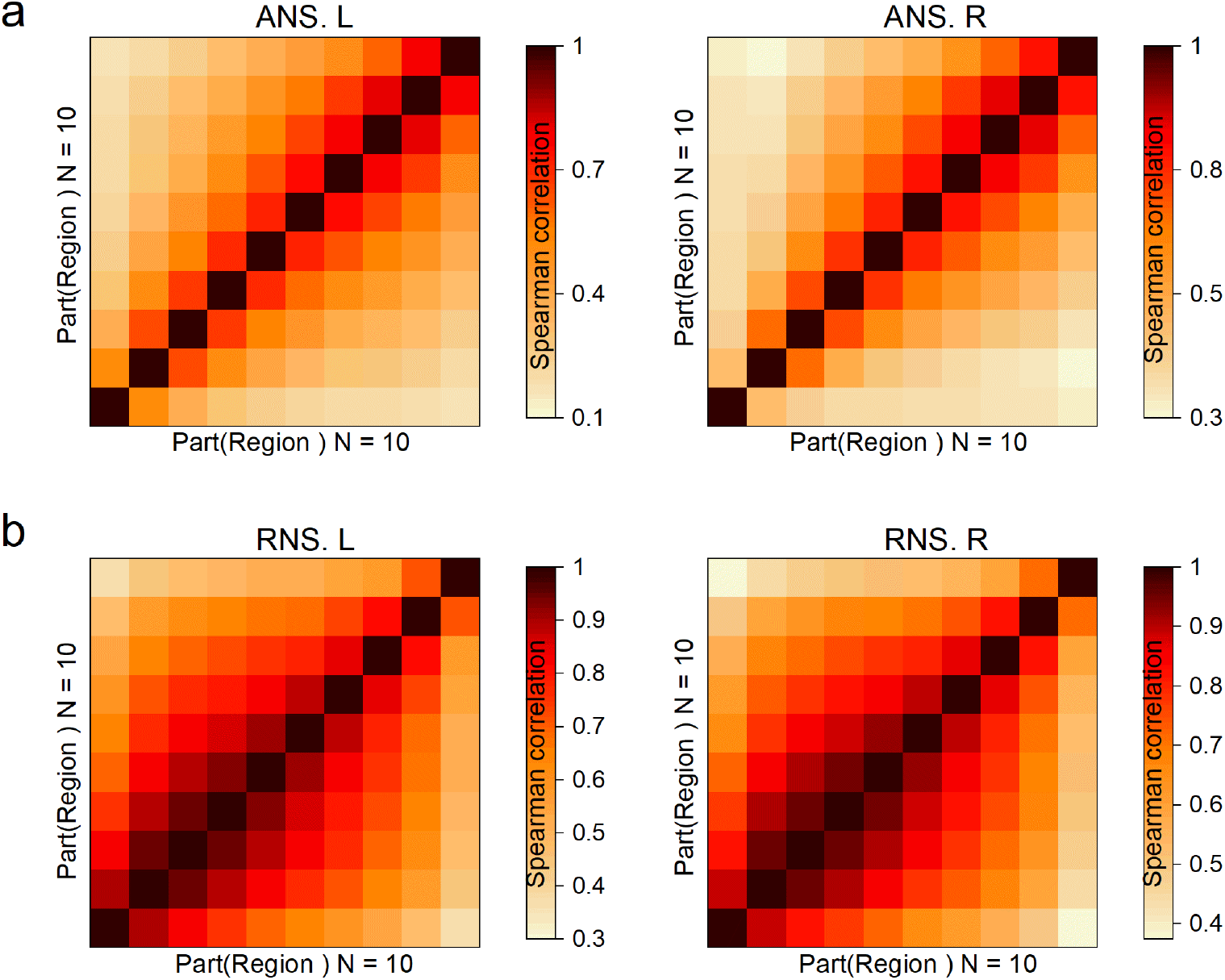
RSA between each pair of regions. We calculated the RDM using the ANS/RNS of voxels in the region. *A_ij_* denotes the similarity between the RDM of region i and the RDM of region j. Spearman correlation analysis was used to calculate similarity.

**Supplementary Fig. 6.**
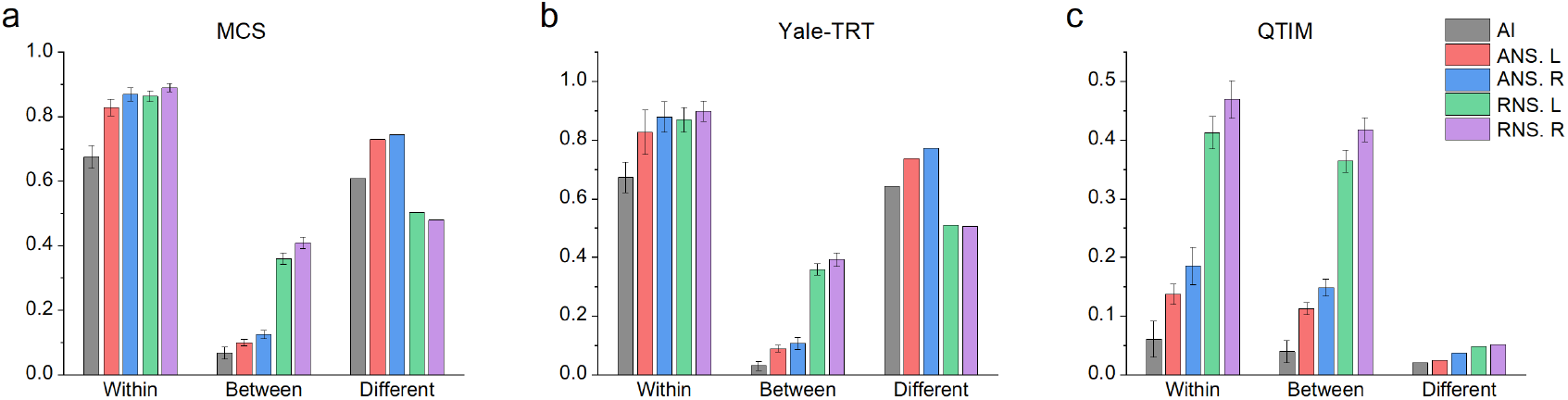
Comparison of ANS and RNS with asymmetric index. Similar to Fig. 2e, f and g, we calculated the stability of the asymmetric index across multiple scans and the heritability of asymmetric index among family members. Two-sample t-test for between-group comparisons. AI: asymmetric index. Error bars indicate SD×1.5

**Supplementary Table 1.** Accuracy of ANS, RNS and asymmetric index (AI) for individual identification and family identification.

| Dataset | AI | ANS |  | RNS |  |
| --- | --- | --- | --- | --- | --- |
|  |  | LH | RH | LH | RH |
| MSC | 100% | 100% | 100% | 100% | 100% |
| RETEST | 100% | 100% | 100% | 100% | 100% |
| QTIM | 27.3% | 77.3% | 59.1% | 86.4% | 86.4% |

**Supplementary Table. 2.** Performance of generated and actual images for gender classification.

| Feature | Accuracy | Precision | Recall | AUC |
| --- | --- | --- | --- | --- |
| Act. L $\oplus$ Act. R | 55.92% | 55.85% | 53.52% | 0.53 |
| Act. L $\oplus$ Recon. R | 54.04% | 53.81% | 50.46% | 0.5 |
| Recon. L $\oplus$ Aat. R | 53.70% | 51.43% | 50.15% | 0.5 |
| Recon. L $\oplus$ Recon. R | 54.26% | 55.42 | 50.70% | 0.5 |

**Supplementary Fig. 7.**
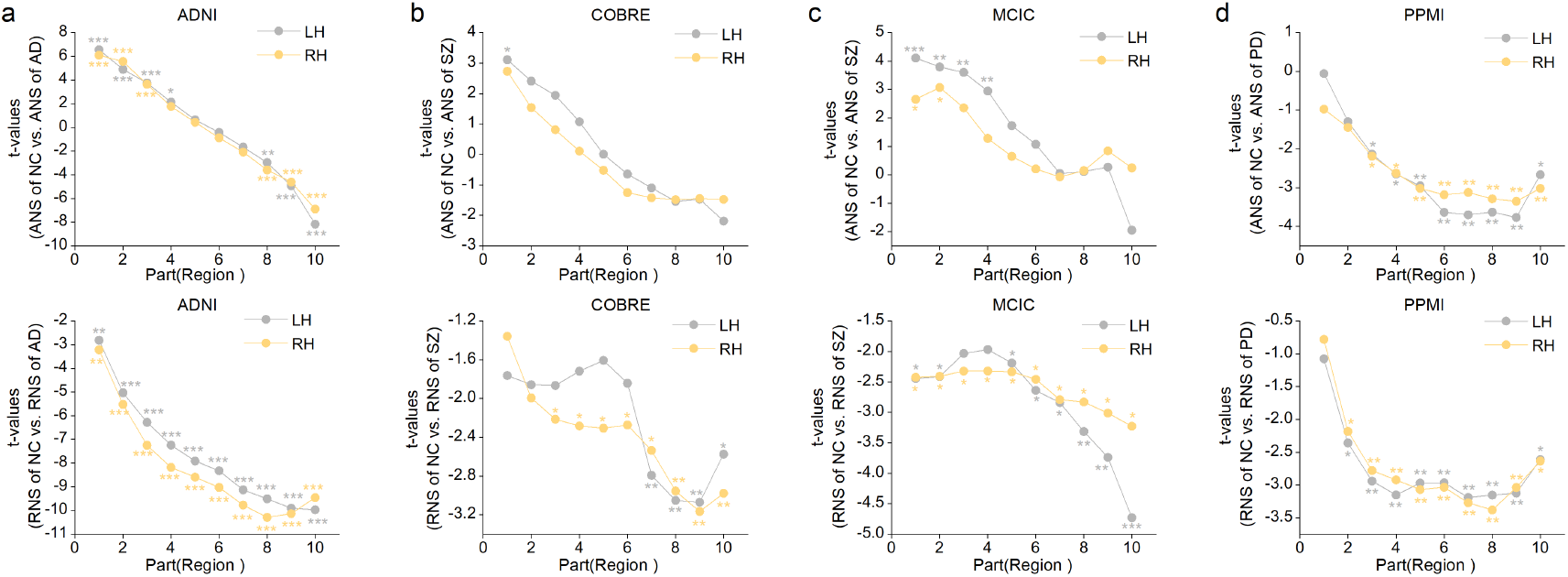
Comparison between the regional ANS/RNS of healthy people and patients. Two-sample t-test for between-group comparisons. The x-axis indicates the index of the region (Supplementary Fig. 4). The y-axis indicates the t-value of the comparison between groups. ***: p < 0.001, **: p < 0.01, *: p< 0.05.

**Supplementary Fig. 8.**
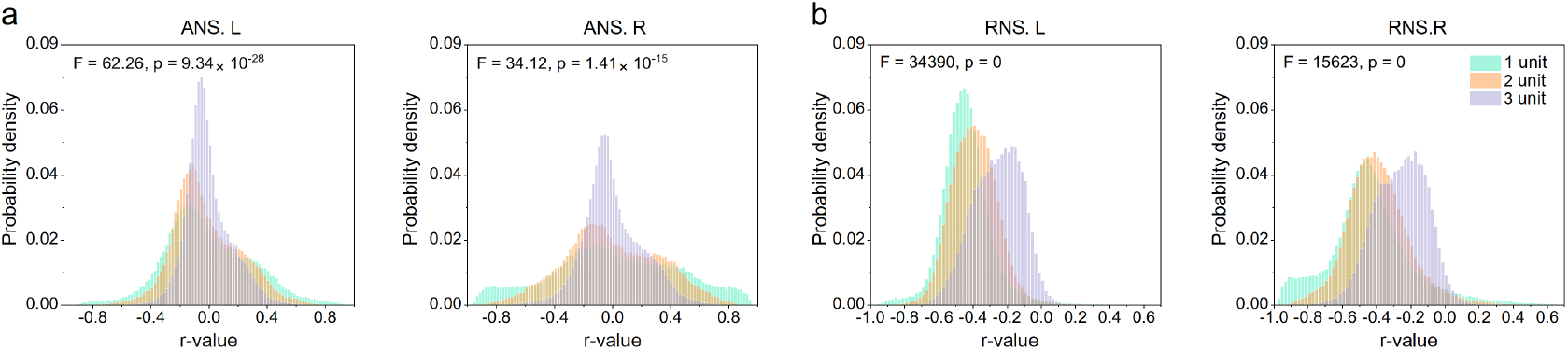
Correlation of the ANS/RNS of each voxel with the GM density of the voxels at a distance K from it. The unit of distance is the length of a voxel (1.5mm). In this study, K = 1, 2 and 3. The X-axis represents the r-value, and the Y-axis represents the probability density. One-way ANOVA was used for comparison between groups.

**Supplementary Fig. 9.**
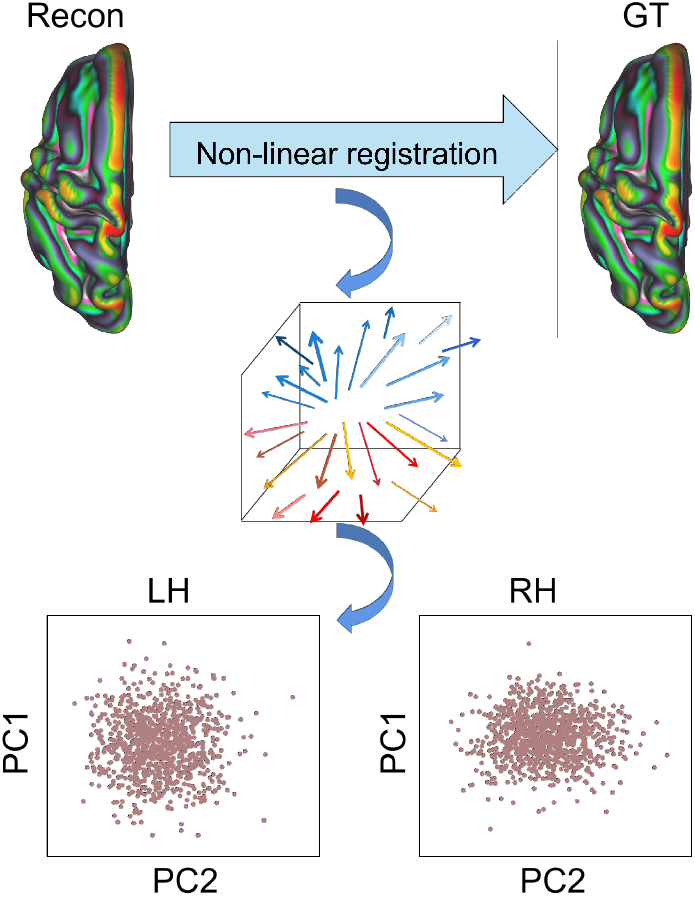
Workflow of deformation-based morphometry analysis.

**Supplementary Fig. 10.**
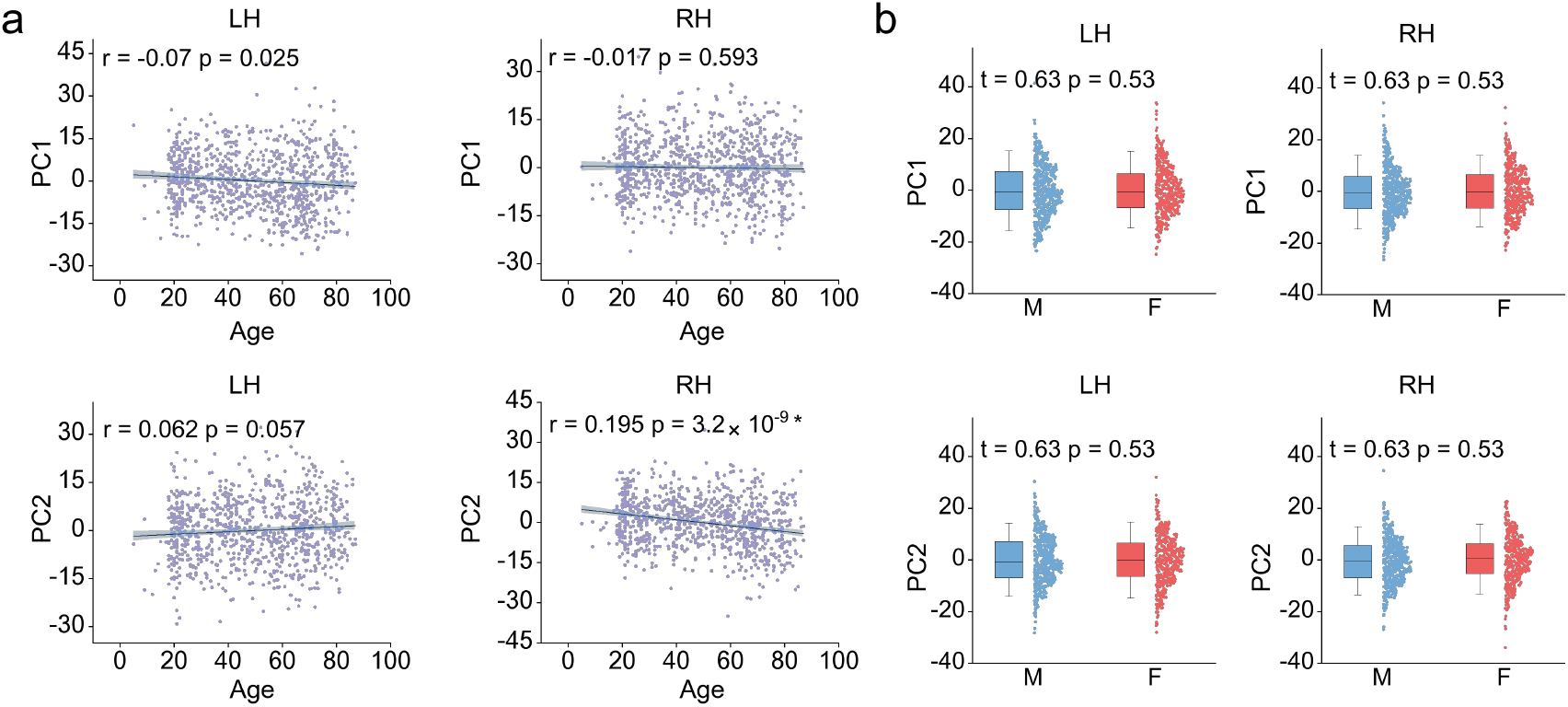
Association between Jacobi map with age and gender. Pearson correlation analysis was used to calculate the similarity between age and PCs of the Jacobi map. The shaded area represents the 95% confidence interval. Two-sample t-test for between-group comparisons. Error bars indicate SD×1.5.

**Supplementary Table 3.** Imaging parameters.

| Dataset | Strength | Repetition Time (ms) | Echo Time (ms) | Flip Angle | FOV (mm) | Voxel size (m<sup>3</sup>) |
| --- | --- | --- | --- | --- | --- | --- |
| IXI | 1.5T or 3T | 9600 | 4.6 | 8° |  |  |
| COBRE | 3T | 2530 | 1.64 | 7° | 256x256 | 1×1×1 |
| Yale-TRT | 3T | 2400 | 1.18 | 8° |  | 1×1×1 |
| Cam-CAN | 3T | 2250 | 2.99 | 9° | 256 × 240 × 192 | 1×1×1 |
| CNP | 3T | 1900 | 2.26 |  | 250 × 250 × 250 | 1×1×1 |
| MCIC | 1.5T or 3T | 2530 | 4.76 | 7° | 256×256×128 | 1×1×1 |
| NUSDAST | 1.5T | 9700 | 4 |  |  | 0.625×0.625×1.5 |
| ADNI | 3T | 2300 | 2.98 | 9° | 256×240 | 1×1×1.5 |
| QTIM | 4T | 1500 | 3.35 |  | 230 | 0.9×0.9×0.9 |
| MCS | 3T | 2400 | 3.74 | 8° |  | 0.8×0.8×0.8 |

The imaging parameters of PPMI are available at http://www.ppmi-info.org. The imaging parameters of Beijing zang, NKI and ICBM are not available.

